# Multiscale Modeling of Vector-Borne Diseases: The Role of Dose-Dependent Transmission

**DOI:** 10.1101/2025.08.22.671686

**Authors:** Fernando Saldaña, Jorge X. Velasco-Hernández, Pauline Ezanno, Hélène Cecilia

## Abstract

The transmission of infectious diseases involves complex interactions across multiple biological scales, from within-host immunological processes to between-host transmission dynamics. While multiscale models have the potential to capture these interactions more accurately, they are often hindered by increased complexity and limited data availability. In this study, we develop a multiscale epidemic model linking host-vector population-level transmission dynamics to within-host and within-vector pathogen dynamics. Our model captures key features of within-vector viral progression and allows bidirectional coupling between within-host and between-host processes. The scales are linked under the assumption of dose-dependent transmission, with the functional form informed by empirical viremia–infectiousness data from arbovirus transmission. Focusing on Dengue, Zika, and West Nile viruses as case studies, we assess how different functional forms of the coupling affect the number of equilibria of the epidemic model. We find that when the transmission is modeled using linear coupling functions, the multiscale model yields the same bifurcation structure of the simpler, uncoupled model, indicating that the linking of scales does not alter the range of possible long-term epidemiological states in such cases. However, nonlinear coupling can induce complex behaviors such as multiple endemic equilibria and backward bifurcations, which the uncoupled model does not capture. These results underscore the importance of carefully selecting coupling functions and provide guidance on when multiscale modeling is essential for understanding and managing vector-borne diseases.

## 1 Introduction

Vector-borne diseases account for roughly 17% of the global burden of communicable diseases [75], with mosquito-borne illnesses posing the greatest threat due to their widespread reach, high transmission rates, and severe health impacts. Mosquitoes involved in disease transmission are responsible for more than 700,000 deaths annually, making them the deadliest animals to humans [29]. Malaria alone accounts for about 85% of these deaths. Dengue virus (DENV) also imposes a major health burden, causing an estimated 96 million symptomatic cases and 40,000 deaths each year [75]. Other significant mosquito-borne diseases include Zika virus (ZIKV), chikungunya virus (CHIKV), yellow fever virus (YFV), and West Nile virus (WNV). This substantial disease burden has driven the development of extensive modeling efforts to understand transmission dynamics and inform control strategies for mosquito-borne diseases (see [60, 65] for reviews).

Infectious diseases, especially zoonotic and vector-borne diseases, spread in a complex manner involving multiple hosts and various temporal and spatial scales. These scales can be broadly grouped into two categories. The between-host scale involves disease dynamics across a host population and can be monitored through incidence and prevalence, whereas the within-host scale encompasses the interactions between a pathogen and a host’s cellular population, including immune response. Despite modelers’ growing interest in linking these scales [18, 56, 21, 31, 36, 41], epidemiological and immunological processes are often studied separately. At the between-host level, Kermack–McKendrick-type models, such as the classical *SIR* (Susceptible-Infected-Recovered) model and its many extensions, are widely used to capture the fundamental dynamics of infectious disease spread e.g. [47, 54, 62]. These models can be tailored to represent scales ranging from localized settings spanning a few meters to large-scale distributions of infections across countries [21, 23]. At the within-host level, a common framework is Perelson’s model [59, 9], which tracks the dynamics of uninfected target cells, infected cells, and free virus particles. This model has been extended to include key components such as innate and adaptive immune responses, as well as antiviral treatment effects (for a review on within-host models see [19]). The within-host scale itself encompasses several biological sub-scales, ranging from molecular and cellular levels to tissue and organ systems [21].

Both theoretical and empirical studies now indicate that immunological and viral dynamics within the host do play a role in shaping the spread of infectious diseases among populations [4, 16, 21, 25, 26, 27, 36, 40, 56]. Gilchrist et al. [34, 33] were among the first to link within-host and between-host dynamics through a mechanistic model, providing a more integrated view of host–parasite coevolution. Specifically, they coupled Perelson’s within-host model with the *SI* epidemic model, under the assumption that the internal state of the host influences both transmission and virulence [33]. Since then, a common strategy for linking within-host and between-host scales has been to express transmission, virulence, recovery, and other epidemiological parameters as functions of within-host variables [34, 33, 56]. This has led to the development of *nested* multiscale models, where the within-host dynamics influence population-level transmission, but not the other way around. More recent work has focused on developing *embedded* multiscale models, which allow for two-way feedback between scales. For instance, Feng et al. introduced bidirectional feedback in multiscale models for environmentally-driven diseases linking pathogen load in the environment to both transmission and within-host dynamics [25, 26, 27]. However, integrating epidemiological factors into within-host processes remains a significant challenge [21, 36]. Moreover, even for diseases transmitted directly between hosts, coupling within-host and between-host models introduces substantial mathematical complexity [3, 2, 17, 34, 55, 58, 64, 72, 76]. The addition of vector dynamics results in two dynamical systems at each level, which significantly increases the dimensionality and nonlinearity of the model, making analytical tractability even more difficult. As a result, most existing multiscale models focus on directly transmitted infections (see [21] for a recent review), leaving vector-borne diseases under-explored. To our knowledge, only a limited number of mechanistic multiscale models using ordinary differential equations (ODEs) have been developed for vector-borne diseases, and these have primarily focused on malaria [1, 32], a parasite which differs substantially from mosquito-borne viral diseases.

A crucial but often overlooked aspect of vector-borne diseases in mechanistic epidemic models is intra-vector viral dynamics (IVD) [38, 51, 60]. IVD describes the virus progression through three key barriers within the vector: (i) the infection barrier, where the virus enters the intestinal epithelium; (ii) the dissemination barrier, where the virus exits the intestinal tract, enters the circulatory system and spreads throughout the vector body; and (iii) the transmission barrier, where the virus is released in the mosquito saliva, enabling transmission to a new host [6]. The time needed to complete this process is known as the extrinsic incubation period (EIP). Once the vector becomes infected, the virus typically remains in its system for the rest of its life, although its concentration in tissues outside the salivary glands may decrease with time [48]. In the case of DENV, variations in IVD have been shown to influence the likelihood and scale of human dengue outbreaks [30]. This highlights the importance of accurately modeling IVD in epidemiological studies.

In this paper, we are interested in developing a multiscale epidemic model linking host-vector population-level transmission dynamics to within-host and within-vector pathogen dynamics. The framework focuses on a single vertebrate host and mosquito vector species but is designed to be flexible, allowing adaptation to a range of mosquito-borne diseases through appropriate parameter calibration. For parsimony and analytical tractability, the multiscale model is formulated using deterministic subsystems based on ODEs. A distinctive aspect of our framework is its integration of within-host and between-host dynamics in a vertebrate–vector system, achieved through the explicit modeling of viral processes inside the vector. Following the modeling paradigm of Gilchrist et al. [34, 33], we assume that the population-level transmission rates (host-to-vector and vector-to-host) depend on the within-host and within-vector viral loads. The functional form of this relationship remains uncertain for most pathogens [4, 16, 40, 42, 36], leading many models to adopt a simple linear relationship. For arthropod-borne virus (arboviruses), however, substantial empirical evidence suggests that transmission is dose-dependent and often follows nonlinear patterns [4, 16, 42, 49], at least for host-to-vector transmission. In this work, we incorporate a range of nonlinear transmission functions based on empirical viremia-infectiousness data observed in DENV, ZIKV, and WNV infections improving the biological relevance of the model. These functions ensure that the between-host scale is influenced by the within-host scale leading to a nested multiscale model. To achieve an embedded multiscale model, we go further and extend the multiscale approach of Feng et al. [25, 26, 27] to vector-borne diseases, so that the prevalence of the infection in the host and vector populations influence within-vector and within-host dynamics.

Our primary objective is to examine how the functional form of the coupling influences the qualitative behavior and long-term dynamics of the system. Consequently, the simulation results based on specific parameter values are not intended as precise predictions of disease spread, but rather as illustrative examples. The remainder of this paper is structured as follows. In the next section, we describe the formulation for each of the subsystems composing the multiscale model. Section 3 introduces the dose–response relationships, which characterize the host’s potential to transmit the virus to vectors as a function of viral load. Section 4 presents a rigorous mathematical analysis of the model, supported by relevant numerical simulations. A comprehensive discussion of the findings is provided in Section 5.

## 2 Formulation of the multiscale model

### 2.1 The between-host model

The population-level model describes the spread of the virus within the total populations of vertebrate hosts and mosquitoes, denoted by *N*_*H*_ and *N*_*M*_, respectively. Since our primary focus is on arboviruses transmitted by mosquitoes, we will use the terms “vector” and “mosquito” interchangeably. The mosquito-host structure comprises seven mutually exclusive classes: susceptible hosts (*S*_*H*_), exposed hosts (*E*_*H*_), infectious hosts (*I*_*H*_), recovered hosts (*R*_*H*_), susceptible mosquitoes (*S*_*M*_), exposed mosquitoes (*E*_*M*_), and infectious mosquitoes (*I*_*M*_). Hence, *N*_*H*_ = *S*_*H*_ + *E*_*H*_ + *I*_*H*_ + *R*_*H*_, and *N*_*M*_ = *S*_*M*_ + *E*_*M*_ + *I*_*M*_. The following system describes the mosquito-to-host transmission model:

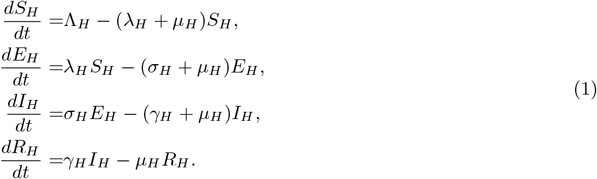

The population of susceptible hosts increases at a constant rate, denoted as Λ_*H*_. Following a mean intrinsic incubation period of 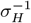, exposed hosts *E*_*H*_ transition into the infectious category *I*_*H*_. Infectious hosts recover at rate *γ*_*H*_, which grants them lifelong immunity. All classes experience a decline due to natural death at rate *µ*_*H*_. Transmission is modeled using a host frequency-dependent force of infection (FoI):

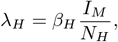

where *β*_*H*_ is the transmission rate from mosquito to host. The following system of ODEs gives the host-to-mosquito transmission model:

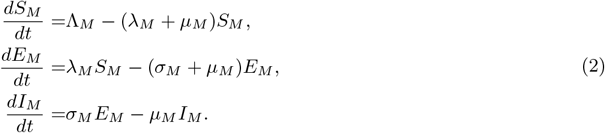

Mosquitoes enter the susceptible class at a constant rate, denoted as Λ_*M*_, and experience natural death at rate *µ*_*M*_. The parameter 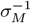 indicates the average extrinsic incubation period. The FoI *λ*_*M*_ is given by

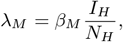

where *β*_*M*_ is the transmission rate from host to mosquito. The FoIs *λ*_*H*_ and *λ*_*M*_ imply that the rate at which a mosquito bites hosts remains unchanged across host densities. In other words, the mosquito does not bite more often because more hosts are around. However, the number of bites that a host receives is proportional to the current mosquito-to-host ratio. Consequently, if the host population is held constant, an increase in mosquito density leads to a higher biting rate per host. This host frequency-dependent FoI has been used extensively to model transmission in several vector-borne diseases, improving over the mass action and susceptible frequency-dependent FoIs (see [74, 15] and the references therein). Importantly, under the assumptions of a constant host population and a fixed mosquito biting rate, this formulation is mathematically equivalent to the mosquito-to-host transmission term in the classical Ross-Macdonald model [65].

### 2.2 The within-host model

We describe the dynamics of arbovirus infection within a host using a simple mechanistic ODE model with an eclipse phase [19]. The model equations are as follows:

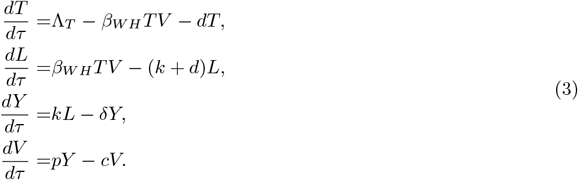

The variable *τ* represents the time scale at which the dynamics within the host occur and could differ from the population-level time scale *t* (see Section 4). Susceptible target cells (*T*) are generated at a constant rate Λ_*T*_, while these cells have a natural death rate of *d*. Viral particles (*V*) infect susceptible target cells with a constant transmission rate *β*_*WH*_ following the mass action law. Newly infected cells spend time in an eclipse phase (*L*) before they become productively infected cells (*Y*) at rate *k* or are cleared at rate *d*. Productively infected cells release the virus at rate *p* and are removed at rate *δ* ≥ *d*. Virus particles are removed at rate *c*. The model (3) does not explicitly account for the effects of the immune response; however, these effects are implicitly incorporated through the virus clearance rate and the death rate of infected cells, which is equivalent to assuming a constant effect of the immune response [7, 8]. This system captures viral dynamics using a mean-field approach, representing the average dynamics within a typical host.

### 2.3 The within-vector model

Following [67], we consider a minimalistic within-mosquito model that follows the virus in the midgut (*W*_*m*_) and in the salivary glands (*W*_*s*_):

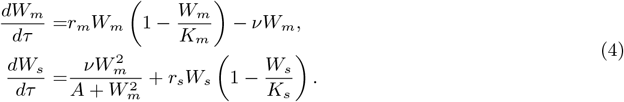

Here, *r*_*m*_ and *K*_*m*_ represent the virus growth rate and carrying capacity in the midgut, respectively. The first term on the right-hand side of the equation for *W*_*m*_ represents the logistic growth of the virus in the midgut. Similarly, *r*_*s*_ and *K*_*s*_ represent the virus growth rate and carrying capacity in the salivary glands of mosquitoes. The parameter *ν* represents the rate at which the virus spreads from the midgut to the salivary glands. The time the virus takes to develop inside a vector before reaching the salivary glands is modeled by a Hill function with a coefficient 2. The Hill function decreases the virus entry into the salivary glands when *W*_*m*_ is low compared to *A*, meaning that higher values of *A* simulate a longer EIP [55]. The EIP is known to vary depending on the pathogen, vector species, temperature, and other environmental factors [39]. Furthermore, the EIP is significant compared to mosquito lifespan; hence, transmission varies according to the relationship between both periods [67, 51]. For example, a shorter EIP means mosquitoes become infectious faster, increasing the chances of successfully transmitting the virus, potentially multiple times, before they die [11].

### 2.4 The coupled multiscale model

Now, we propose an explicit link between the population-level transmission model (1)-(2) and the within-host and within-vector subsystems (3)-(4). The classical approach to incorporate within-host dynamics into the population-level epidemiological process is to assume that there is a functional relationship between pathogen transmission rates and within-host state variables, as demonstrated in the work of Gilchrist et al. [34, 33]. We follow this approach and assume that the transmission rate from mosquito to host *β*_*H*_ (*W*_*s*_) depends on the viral load in the mosquito’s salivary glands *W*_*s*_ [51]. Likewise, we assume that the transmission rate from host to mosquito *β*_*M*_ (*V*) depends on the concentration of free virions within a host *V*. Although it is widely recognized that the pathogen load within the host should influence transmission rates, the exact form of the dose-response relationship remains poorly understood for most diseases [4, 16, 40, 42, 36]. At this stage, we only assume that the functions *β*_*H*_ (*W*_*s*_) and *β*_*M*_ (*V*) are nonnegative, continuous, and satisfy the conditions *β*_*H*_ (0) = *β*_*M*_ (0) = 0. The specific functional relationships between disease transmission rates and within-host/vector variables will be addressed in the subsequent section.

Most models aiming to integrate immunological and epidemiological dynamics face the challenge of determining the most suitable methods for incorporating epidemiological factors into cellular processes at the micro scale [21, 36]. As a result, in many studies, the between-host scale is influenced by the within-host scale but not vice-versa. Here, aiming to obtain bidirectional feedback between the within-host and between-host scales, it is assumed that the abundance of infected mosquitoes measured by *I*_*M*_ */N*_*M*_ := *i*_*M*_ increases the within-host viral load via a feedback function *g*_*H*_ (*i*_*M*_) where the function *g*_*H*_ has the following properties: 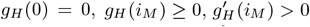, and 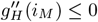 [26, 25, 27]. It is important to remark that while this approximation is convenient for a mean-field ODE model, it overlooks the relatively brief interaction between the within-host and the between-host dynamics and the fact that the continuity assumption described above can be generalized to more realistic interaction functions. The effect of the infected host population on within-mosquito virus dynamics is captured by a function *g*_*M*_ (*i*_*H*_), where *I*_*H*_ */N*_*H*_ := *i*_*H*_ is the proportion of infected hosts. Since the within-mosquito model (4) does not account for the full mosquito life cycle and instead tracks the virus only in the midgut and salivary glands, we assume that *g*_*M*_ has two main effects. A fraction *q* increases the viral load in the midgut, speeding up virus dissemination within the mosquito. The remaining fraction 1 − *q* boosts the carrying capacity of the salivary glands, increasing the ability of mosquitoes to transmit the virus. As with *g*_*H*_, function *g*_*M*_ satisfies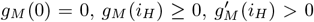, and 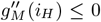 [26, 25, 27]. Under these assumptions, the full multiscale model is given by the following system of ODEs:

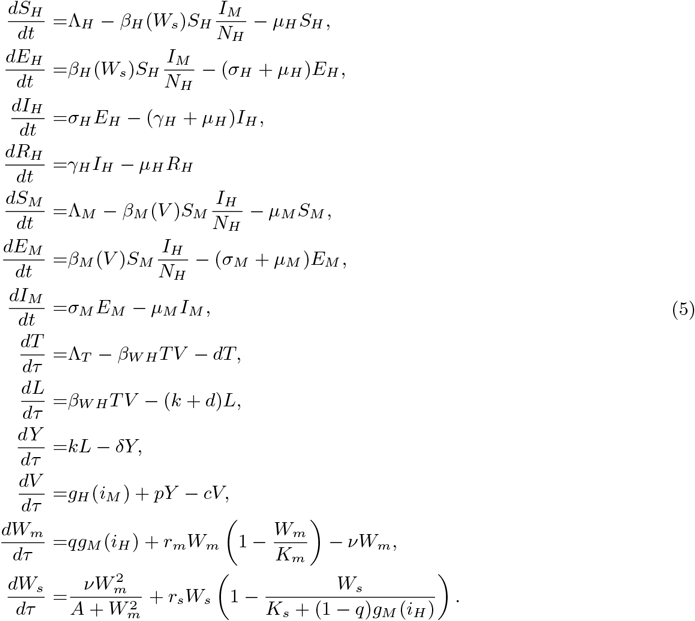

The parameters of the multiscale model (5), together with initial conditions, units and average values, are described in section S1 of the Supplementary Material (SM) and summarized in Supplemental Tables S1-S3. For illustrative purposes and based on available data, we compiled parameter values for both the between-host and within-host subsystems for three major arboviruses: DENV, ZIKV, and WNV. For the within-vector model, however, parameter estimates were only available for ZIKV-infected mosquitoes.

## 3 Dose-response relationships in arbovirus transmission

Most mathematical models exploring arbovirus transmission dynamics at the population level do not explicitly incorporate within-host or within-vector viral load. Instead, they typically rely on threshold-based assumptions, classifying vertebrate hosts as either infectious or non-infectious to vectors based on an implicit viral load cutoff [53]. Similarly, once vectors are categorized as infectious, these models assume that all mosquitoes transmit similarly to hosts. However, growing evidence suggests that arbovirus transmission is dose-dependent, with viral load influencing the probability of successful transmission events [4, 16, 42, 49].

After mosquitoes are exposed to viremic blood, either through an artificial blood meal or a live host, the presence of the virus within mosquitoes can be assessed by analyzing distinct anatomical compartments. These include: (i) the body (commonly referring to the abdomen, excluding head, legs, and wings), ii) the legs or the head, and iii) the salivary glands or the saliva itself. These analyses help elucidate whether the virus has successfully overcome the infection, dissemination, and transmission barriers within the mosquito [6]. As these measurements require the mosquito to be killed, they can only be conducted at a single time post feeding. Consequently, estimating the dose-response relationship between host viremia and mosquito infection probability necessitates sacrificing separate cohorts of mosquitoes, each exposed to different initial viral doses.

Estimating the EIP further multiplies the number of mosquito batches sacrificed at distinct times post feeding for a given initial dose.

We compiled published dose-response data linking infectious titers in vertebrate hosts to the probability of mosquito infection for DENV, ZIKV, and WNV (see Figure 1). All data were drawn from studies where mosquitoes fed directly on live hosts, which more accurately reflects natural transmission conditions. This is important because mosquitoes generally require higher viral doses to become infected via artificial blood meals [61, 52].

**Figure 1:**
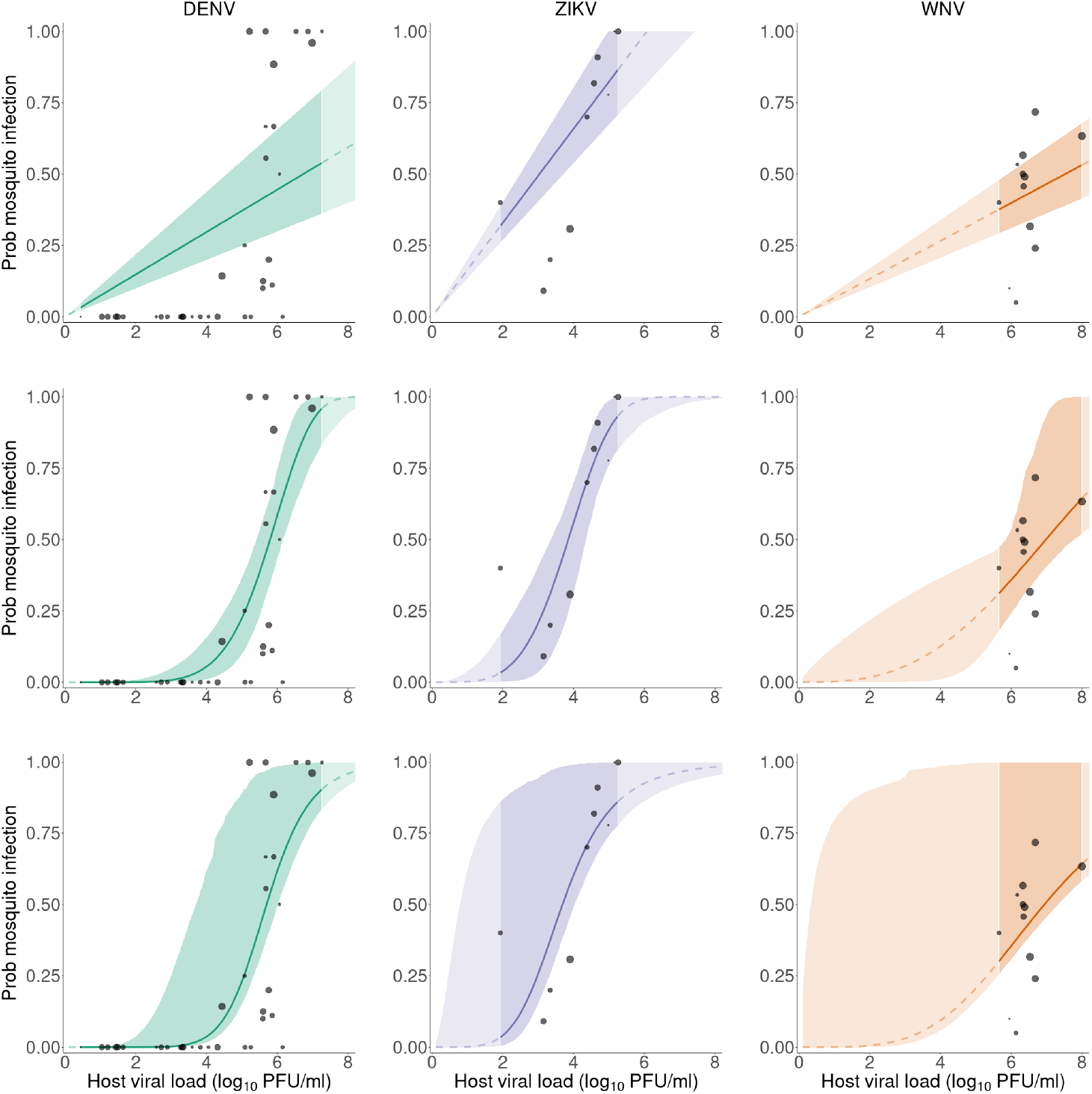
Dose–response relationships between vertebrate host infectious titers and the probability of mosquito infection. Points show data for DENV (first column), ZIKV (second column), and WNV (third column), sourced from [22], [42], and [69], respectively. Point size is proportional to the number of mosquitoes tested in the batch, but the scale is virus-specific ([1;26] for DENV, [9;13] for ZIKV, [10;60] for WNV). The three rows display model fits using a linear response (top), the Ferguson functional response (middle), and the Hill function (bottom). Curves represents the best fit and shaded areas correspond to 95% confidence intervals. Darker color and solid line used within the data range, lighter color and dashed line used outside the data range. All fits were obtained using a betabinomial likelihood. Note that mosquito infection was measured in mosquito legs for DENV and ZIKV, and in bodies for WNV, which makes them not directly comparable.

For DENV, we used data from a study by Duong et al. [22], conducted in Cambodia during the 2012 and 2013 summer seasons (June–October). Mosquito infection was assessed by detecting virus in legs and wings after a 13–16 day EIP. The experiments used *Aedes aegypti* mosquitoes, and human viremia was quantified by qRT-PCR in genome copies. We converted these values to infectious titers (plaque forming units, PFU) using a conversion factor from [10]. Although the dataset includes dose-response data for DENV serotypes 1, 2, and 4, we focused exclusively on DENV-4 to avoid cases where transmission to mosquitoes happened with undetectable viremia (which happened for the other serotypes).

For ZIKV, we used data from experimental infections of non-human primates, specifically the data for cynomolgus macaques (*Macaca fascicularis*) described in [42]. The animals were infected through mosquito bites with a sylvatic ZIKV strain (DakAR41525), and infectious viremia was measured in PFU. *Aedes albopictus* mosquitoes were used in the study, and infection was determined through the presence of virus in the legs after a 14-day EIP.

For WNV, we used data from Vaughan et al. [69], which tracked the relationship between viremia in common grackles (*Quiscalus quiscula*) and subsequent infection of *Culex pipiens* mosquitoes. We selected the data for the Area B mosquito strain, as it was a more recent colony. Mosquitoes were assayed individually for WNV presence using plaque assays conducted 3 to 7 days post-feeding with infection assessed by detecting the virus in mosquito bodies. We note that some birds in this experiment were also co-infected by microfilarial parasites, but this did not affect WNV infectivity to mosquitoes.

To explore the dependence of mosquito infection rates on host viral titers, we propose the following functional forms

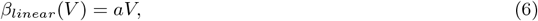

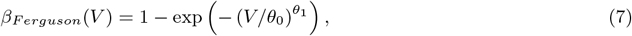

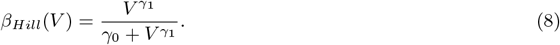

In the functional forms (6)-(8), the variable *V* represents the host infectious titer measured in log_10_ PFU/ml. The parameters *a, θ*_0_, *θ*_1_, *γ*_0_, and *γ*_1_ are all positive constants that will be estimated from the data. Equation (6) describes a linear function between the host viremia and the probability of mosquito infection. Equation (7) is a sigmoidal functional form that was previously used in DENV modeling [28]. Finally, (8) defines a Hill function which is commonly used to represent sigmoidal biological responses.

For each dataset, we fitted the three functional forms defined in equations (6)–(8), testing both a binomial or a betabinomial likelihood, the latter accounting for overdispersion in the data. The fitting procedure is detailed in section S2 of the SM. Model selection was based on the corrected Akaike Information Criterion (AICc), with results summarized in Supplemental Table S4. Point estimates and corresponding 95% confidence intervals for estimated parameter are reported in Supplemental Table S5.

Figure 1 shows the fitted curves for each functional form and virus, based on the betabinomial likelihood, which consistently outperformed the binomial likelihood (Table S4). Model selection showed that the linear model (6) best described the ZIKV and WNV data, while for DENV, the Ferguson and Hill models provided equally good fits, with identical AICc values (Table S4). For ZIKV, the difference in AICc value between the linear and the Ferguson fits was below 2 points.

The estimated dose–response relationships quantify the link between pathogen load and infectiousness, providing critical insight for bridging within-host dynamics with population-scale transmission models. In this context, the fitted functional forms serve as proxies for the host-to-mosquito transmission rate, *β*_*M*_ (*V*). A similar dose-response phenomenon is expected to take place for vector-to-host transmission, but precise estimates for this process are lacking for most arboviruses [4]. Typically, researchers determine the infectious dose by injecting hosts with serial dilutions of the virus and then monitoring for infection. However, this method does not accurately replicate natural transmission, as mosquito saliva contains proteins that modulate both viral infectivity and the host’s immune response [63, 37, 71]. Although experimental infections via mosquito bites are feasible [12, 42], they are technically challenging, costly, and do not permit precise control or measurement of the virus dose delivered [4]. Even when virus quantification is attempted through forced salivation following a blood meal, the measured amount may not accurately represent what was actually transmitted to the host [35, 57]. Given these limitations, we chose to use the same dose–response functional forms for both host-to-vector and vector-to-host transmission in our multiscale modeling framework, acknowledging this simplification while enabling tractable analysis.

## 4 Model analysis

Time-scale separation methods are crucial for simplifying the analysis and simulation of multiscale epidemic models [27, 50, 72, 76]. A standard method for handling time-scale separation involves treating the fast and slow variables independently using a perturbation expansion, where a small parameter is introduced. This approach is biologically relevant because both the within-host and within-vector dynamics typically operate on a faster time scale compared to population-level between-host dynamics. Under these assumptions, the multiscale model is usually divided into a “fast” subsystem (representing within-host processes) that governs short-term dynamics and a “slow” subsystem (the population-level transmission model) that captures long-term dynamics. We follow this approach assuming that the slow (*t*) and fast (*τ*) time scales are related as *t* = *ϵτ* where 0 *< ϵ* ≪ 1.

### 4.1 The fast system

To investigate the fast dynamics, we need to express the between-host subsystem in terms of the within-host time scale *τ* and then assume that *ϵ* → 0. This implies that the derivative of the vector of slow state variables *x* := (*S*_*H*_, *E*_*H*_, *I*_*H*_, *R*_*H*_, *S*_*M*_, *E*_*M*_, *I*_*M*_)^*T*^ is equal to zero, see section S3 in the SM. Hence, we can investigate the within-host subsystems, assuming the slow state variables are constant.

#### 4.1.1 Dynamics of the within-host model

In the absence of feedback from the between-host system, that is, with *g*_*H*_ (*i*_*M*_) = 0, the within-host model (3) has a virus-free equilibrium denoted 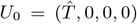 where 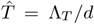 represents the total number of target cells without infection. The basic reproduction number determines the stability properties of *U*_0_. In a within-host context, this number represents the average number of second-generation infectious cells generated by a single infected cell in a population where all cells are initially susceptible [19]. We derive the within-host basic reproduction number, denoted *R*_*w*_, for model (3) using the next-generation operator [20] and the method of van den Driessche and Watmough [68] yielding (see section S4 of the SM)

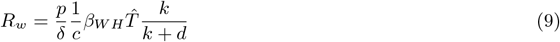

The expression (9) for *R*_*w*_ can be interpreted as follows. The burst size *p/δ* gives the number of new viral particles produced and released during the lifetime of a productively infected host cell. The average time a viral particle remains infectious is 1*/c*. During this period, infectious virions can infect an average of 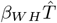 target cells in a fully susceptible cell population (at the viral-free equilibrium). However, only a fraction *k/*(*k* + *d*) of these cells will survive the eclipse phase and become productively infected cells.

If the within-host basic reproduction number satisfies *R*_*w*_ *>* 1, system (3) has a unique positive endemic equilibrium

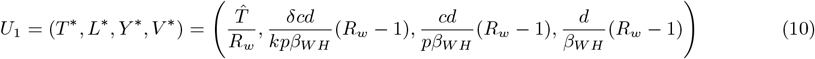

As a consequence of Theorem 2 in [68], the DFE *U*_0_ is locally asymptotically stable (LAS) if *R*_*w*_ *<* 1, and unstable if *R*_*w*_ *>* 1.

Figure 2 presents the viral load measured in log_10_ plaque-forming units per milliliter (PFU/mL) using parameters corresponding to DENV, ZIKV, and WNV. The mean parameter values used in the simulations, along with their sources, are detailed in section S1 of the SM. Under these parameter conditions, the within-host reproduction number satisfies *R*_*w*_ *>* 1, indicating a supercritical regime where the system approaches an equilibrium with a positive viral load but relatively close to zero, resembling an acute infection.

**Figure 2:**
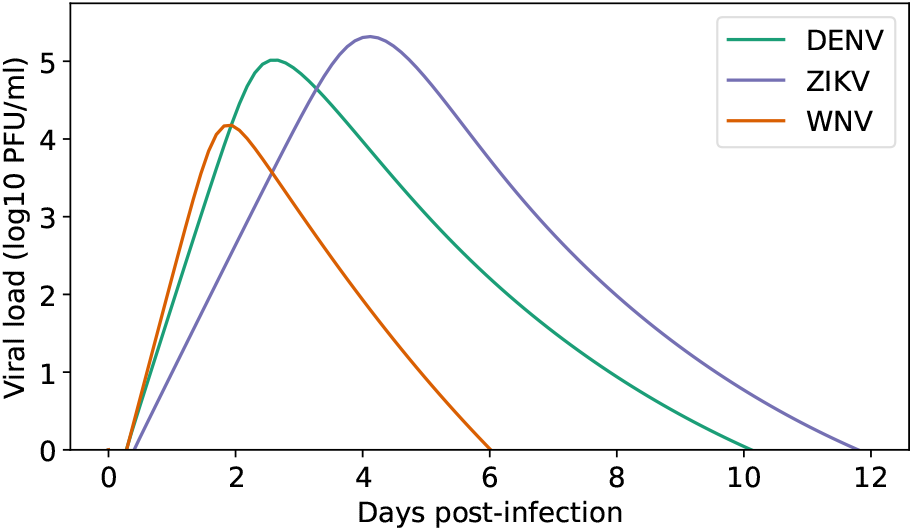
Viral load measured in log_10_ plaque-forming units per milliliter (PFU/ml) given by the within-host model (3) using parameters for DENV, ZIKV, and WNV, respectively. Mean parameter values are given in the Supplemental Table S2.

Parameter estimation in models of viral dynamics is inherently influenced by the availability and reliability of empirical data, often resulting in substantial uncertainty [77]. To systematically evaluate the impact of this uncertainty on model outputs, we performed a global sensitivity analysis (GSA) focused on the equilibrium viral load, *V* ^*^. Specifically, we applied Sobol’s method, a variance-based approach that quantifies the contribution of each input parameter to the overall variance of the model outputs [66]. This method provides both first-order indices, reflecting the isolated effect of individual parameters, and total-order indices, which account for the combined effects of parameters, including all higher-order interactions. Both indices are non-negative, with total-order indices being greater or equal than their corresponding first-order values. The implementation used the open source SALib library [45]. Figure 3a shows that the most influential p arameters f or the equilibrium viral load are similar for the three illustrative pathosystems. Those are the recruitment rate of susceptible target cells Λ_*T*_, the death rate of infected target cells *δ*, the in-host viral production rate per cell *p*, and the in-host clearance rate of free virions *c*.

**Figure 3:**
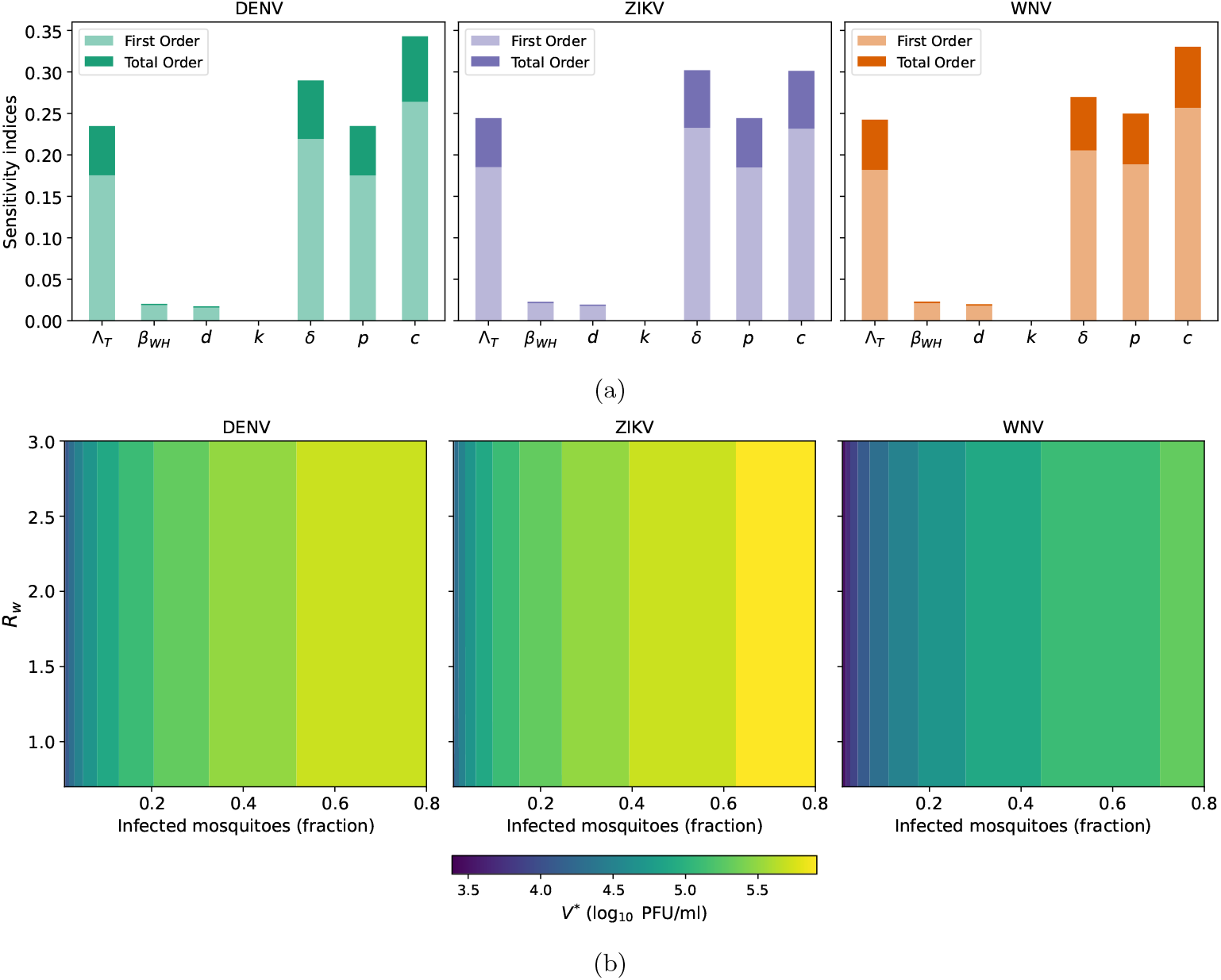
(a) First and total order Sobol sensitivity indices for the within-host equilibrium viral load for DENV, ZIKV, and WNV, respectively. The ranges used for the parameter values in the sensitivity analysis are based on 50% deviation from the mean values presented in Supplemental Table S2. (b) Contour plot of the equilibrium viral load *V* ^*^ in log_10_ scale as a function of the within-host reproduction number *R*_*w*_ (y-axis) with values in the interval [0.8, 3] and *g*_*H*_ (*i*_*M*_) = *ai*_*M*_ (x-axis) with *a* = 10^7^ and *i*_*M*_ ∈ [0.01, 0.8].

Now, we consider the case where there is a feedback from the population-level dynamics. Since the feedback function *g*_*H*_ (*i*_*M*_) does not change with time on the fast time scale *τ*, we can assume *g*_*H*_ (*i*_*M*_) equals a positive constant. Under this assumption, the within-host model no longer has a virus-free equilibrium. To find an equilibrium with a positive viral load

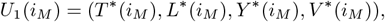

we can assume *V* ^*^(*i*_*M*_) *>* 0 and express the rest of state variables as functions of *V* ^*^(*i*_*M*_) as follows

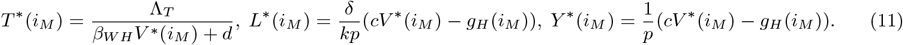

The values of *V* ^*^(*i*_*M*_) are given by the roots of *V* ^2^ − *A*_1_*V* − *A*_2_ = 0, where

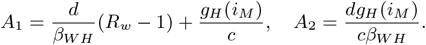

Since *A*_2_ *>* 0, then 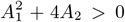 and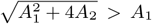. Therefore 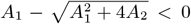 and the only biologically feasible equilibrium solution is

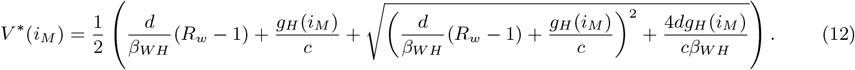

A direct computation shows that the derivative of *V* ^*^(*i*_*M*_) is positive; therefore, this equilibrium viral load is an increasing function of *i*_*M*_ (see also Figure 3b). As *i*_*M*_ → 0, the limiting value of the viral load is

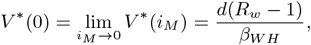

thus *V* ^*^(0) is only biologically relevant for *R*_*w*_ *>* 1. The linearization matrix of system (3) at *U*_1_(*i*_*M*_) is

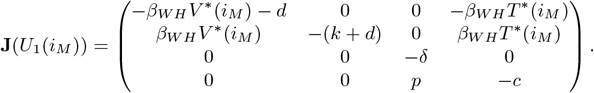

The characteristic polynomial is *P* (*λ*) = *λ*^4^ + *a*_1_*λ*^3^ + *a*_2_*λ*^2^ + *a*_3_*λ* + *a*_4_, where the coefficients are

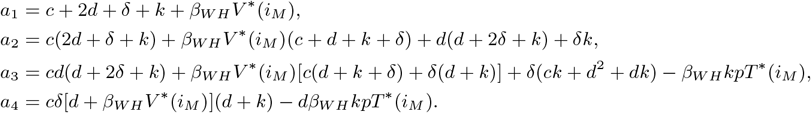

From (11), we have *d* + *β*_*WH*_ *V* ^*^(*i*_*M*_) = Λ_*T*_ */T*^*^(*i*_*M*_), therefore we can rewrite *a*_4_ as

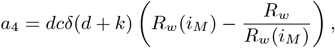

where

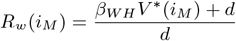

can be interpreted as the within-host reproduction number as a function of *i*_*M*_ *>* 0. Observe that *R*_*w*_ (*i*_*M*_) is an increasing function, *R*_*w*_ (*i*_*M*_) *>* 1 for *i*_*M*_ *>* 0 and *R*_*w*_ (0) = *R*_*w*_. Therefore, if *i*_*M*_ *>* 0 then *a*_4_ *>* 0. Similarly, it can be shown that *i*_*M*_ *>* 0 implies *a*_3_ *>* 0. The coefficients of *P* (−*λ*) are 1, −*a*_1_, *a*_2_, −*a*_3_, *a*_4_. Since the degree of the characteristic polynomial is 4 and the number of sign changes is exactly 4, by Descartes’ rule of signs, all the roots must be negative, which implies the following result:

##### Theorem 1.

*If there is no feedback from the population level dynamics (i*_*M*_ = 0*) and R*_*w*_ *<* 1, *the within-host model* (3) *has a virus-free equilibrium U*_0_ *which is LAS. If i*_*M*_ *>* 0 *the within-host model* (3) *has a unique equilibrium with positive viral load which is LAS*.

The above theorem suggests that, when population-level feedback is considered, the virus can persist within the host, similar to a chronic infection. This is not necessarily the case for most vector-borne diseases. However, we note that the within-host model (3) represents an average host. Therefore, if the disease persists at the population level, then there should be an infected host at each point in time. Moreover, although rare, some arboviruses, such as WNV, can cause chronic infections that periodically reactivate, allowing the virus to survive throughout the seasons, including winter [4, 73]. The relationship between equilibrium viral load *V* ^*^, within-host reproduction number *R*_*w*_, and *g*_*H*_ (*i*_*M*_) for DENV, ZIKV, and WNV is shown in Figure 3b. The variation in the value of *R*_*w*_ is achieved by varying parameter *β*_*WH*_. Figure 3b shows that *V* ^*^(*i*_*M*_) is an increasing function of *i*_*M*_ and in a smaller magnitude, of *R*_*w*_.

#### 4.1.2 Dynamics of the within-vector model

Here, we investigate the stability properties for the within-vector model (4) without feedback from the population-level model, that is, with *g*_*M*_ (*i*_*H*_) = 0. The equation for the virus in the midgut is independent of the equation for the virus in the salivary glands, so we can first analyze the dynamics for *W*_*m*_. Since the equation for *W*_*m*_ is a Bernoulli differential equation, the substitution 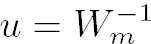 gives a first-order linear differential equation from which we obtain the following solution

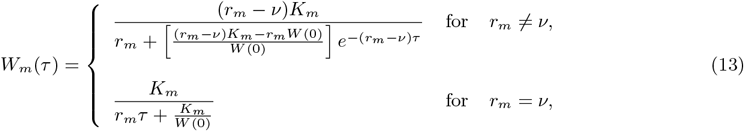

where *W* (0) *>* 0 is a positive initial condition. The following result follows from the explicit solution (13):

##### Theorem 2.

*If r*_*m*_ *> ν and W*_*m*_(0) *>* 0, *then W*_*m*_(*t*) *remains positive for all τ >* 0 *and*

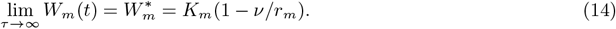

*If r*_*m*_ ≤ *ν, then W*_*m*_(*τ*) *converges to zero as τ* → ∞.

For the equilibrium 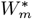 to be positive and stable, it is necessary that viral growth rate in the midgut is larger than the dissemination rate i.e. *r*_*m*_ *> ν*. Once a mosquito gets infected, the virus will typically stay in its system for the rest of its life [48]. Therefore, it is plausible to assume that *r*_*m*_ *> ν*, so (14) holds. In this case, the equilibrium for the virus in the salivary glands 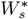 is given by the solution of the next equation

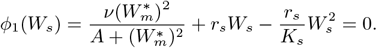

Therefore, the only biologically feasible equilibrium is given by

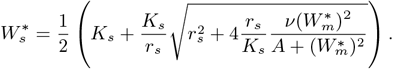

Furthermore,

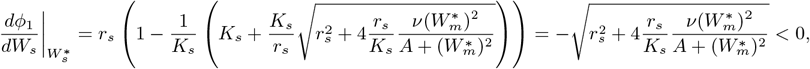

hence 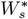 is LAS. This suggests that a virus acquired during a blood meal eventually reaches the salivary glands, making all exposed mosquitoes infectious. A typical trajectory of the within-vector model showing this behavior is given in Figure 4a (dashed lines). Yet, we would like to remark that in reality, the virus may not always be able to cross the infection, dissemination, and transmission barriers [51]. Model (4) can indirectly capture this effect, as for certain values of the dissemination rate *ν* and the half-saturation constant *A* the EIP can be long enough such that the mosquito does not become infectious during its lifetime.

**Figure 4:**
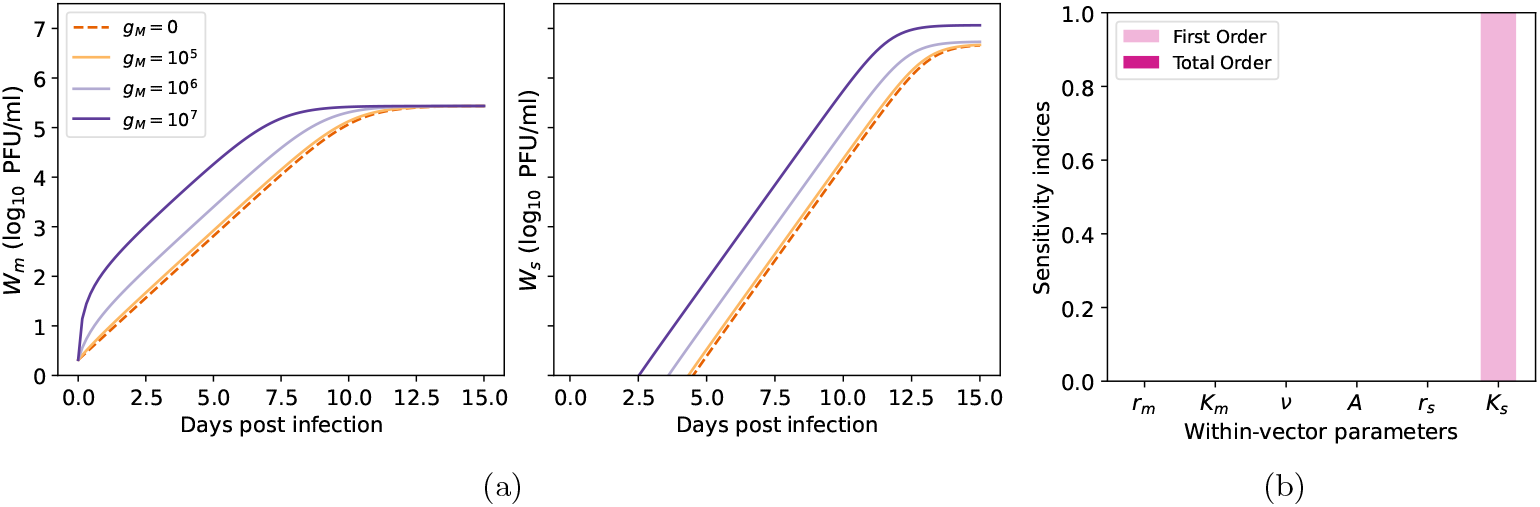
(a) Simulation of the within-vector model for parameter values representative of ZIKV infection given in Supplemental Table S3. The feedback function *g*_*M*_ takes four different constant values 0 (no feedback), 10^5^, 10^6^ and 10^7^ while *q* = 10^−5^. (b) First and total order Sobol sensitivity indices for the equilibrium viral load in the mosquito salivary glands. Observe that the first and total order indices indices are both equal to 1 for parameter *K*_*s*_ and zero for other parameters. The ranges used for the parameter values in the sensitivity analysis are based on 50% deviation from the mean values.

For the within-vector model (4), parameter values were available only for *Aedes aegypti* mosquitoes infected with ZIKV [67]. Consequently, we conducted a global sensitivity analysis to identify the most influential parameters in our within-vector model. Because the mosquito ability to transmit the infection mainly depends on the viral load in its salivary glands, our analysis focused on the equilibrium viral load, 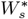, in these glands. The findings indicate (see Figure 4b) that 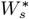 is entirely determined by the salivary gland carrying capacity, *K*_*s*_ (hence the first and total order Sobol sensitivity indices are the same for *K*_*s*_). At the same time, the remaining parameters only affect transient dynamics such as growth, dissemination, and the duration of the EIP.

Let us consider the scenario where the population-level dynamics provide feedback. Again, by time-scale separation arguments, the feedback function *g*_*M*_ (*i*_*H*_) can be considered as a positive constant. Figure 4a displays simulations of the within-vector model using fixed values for *g*_*M*_. Observe that higher values of *g*_*M*_, which lead to a larger viral inoculum, result in more rapid viral growth in the midgut. This accelerated growth promotes faster viral dissemination, allowing the virus to reach greater concentrations in the salivary glands more quickly. Indeed, a larger inoculum shortens the EIP, which is consistent with experimental observations [46]. For long-term dynamics, if *g*_*M*_ (*i*_*H*_) is a positive constant, then 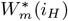 has a unique positive equilibrium solution given by

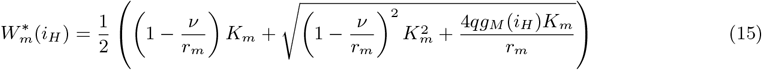

which was obtained as the only positive solution of

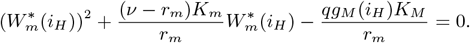

Since 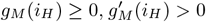, from (15) it follows that 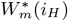 is a monotic increasing function of *i*_*H*_. Further, the derivative of the right-hand side of the equation for *W*_*m*_ in (5) evaluated at 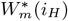 with *g*_*M*_ (*i*_*H*_) *>* 0 is

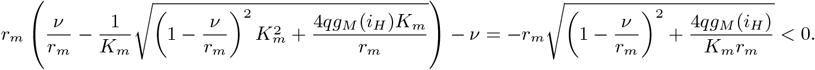

Therefore, if *g*_*M*_ (*i*_*H*_) *>* 0, the level of the virus at the midgut has a unique equilibrium point given by (15), which is LAS. The equilibrium solution for *W*_*s*_ as a function of *g*_*M*_ (*i*_*H*_) is straightforwardly obtained as

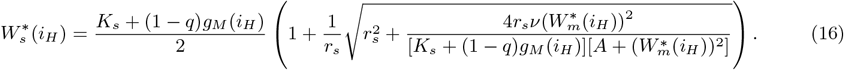

The sign of the derivative of *ϕ*_1_ evaluated at 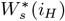 implies that this equilibrium is LAS. Moreover, the equilibrium 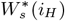 is a monotic increasing function of *i*_*H*_ (see Figure 4a).

### 4.2 The slow system

Next, we examine the dynamics of the between-host system. First, for both the host and the vector, the total population satisfies

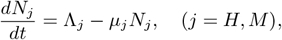

and thus these populations are asymptotically constant to the value 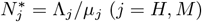. Hence, assuming *N*_*j*_ (0) = Λ_*j*_ */µ*_*j*_, the host and vector population will remain constant at their equilibrium value. Moreover, if *x*_*i*_ = 0, then 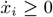 for each of the slow state variables (*i* = 1, 2, …, 7). Therefore, solution trajectories of the between-host system are positively invariant in the region

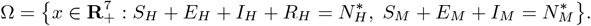

The basic existence, uniqueness, and continuation results for the between-host model hold in Ω, so this system is epidemiologically and mathematically well-posed. Expressing the within-host and within-vector subsystems in terms of the time scale *t* and then assuming that *ϵ* → 0, implies that the within-host subsystems can be treated as being in a steady state on the scale *t* (see section 3 the SM). Under this assumption, the vector of fast state variables *y* := (*T, L, Y, V, W*_*m*_, *W*_*s*_)^*T*^ is at the positive equilibrium and thus the transmission rates are constant 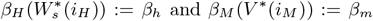. This allows us to study slow dynamics decoupled from the within-host and within-vector processes. We start computing the between-host reproduction number, denoted *R*_*h*_. Using the next-generation method [20, 68], we obtain (see section S4 of the SM):

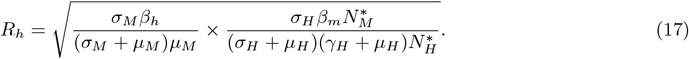

The first term under the square root reflects the number of secondary host infections caused by an infected vector. The second component represents the number of secondary vector infections that result from a typical infected host. The square root in (17) represents the geometrical mean of secondary infections in a single generation. Removing the square root would represent secondary host infections resulting from the introduction of a primary host infection, i.e two generations [20, 68]. The reproduction number (17) is based on a host frequency-dependent force of infection, and hence it depends on the vector-to-host ratio. Reducing the vector population lowers *R*_*h*_, while reducing the host population increases *R*_*h*_ (see Figure 5). Though this may seem counterintuitive, it reflects the assumption that vector biting rates are independent of host density. As a result, when the host population decreases, the remaining hosts are bitten more frequently [74, 15].

**Figure 5:**
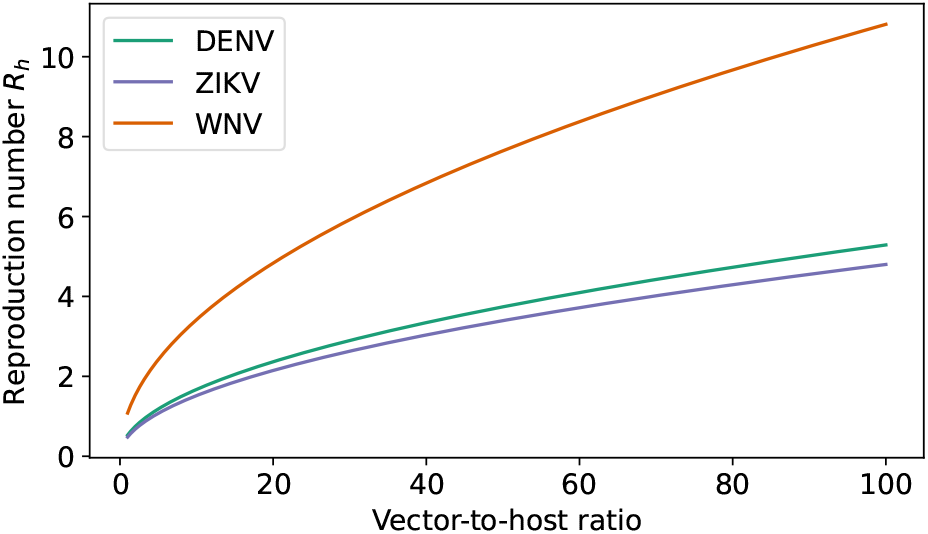
Between-host reproduction number *R*_*h*_ as a function of the vector-to-host ratio 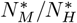. Mean parameter values are given in Supplemental Table S1.

A simple computation shows that the uncoupled between-host model has a unique disease-free equilibrium 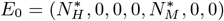.To obtain the endemic equilibrium, it is convenient to define the following auxiliary constants:

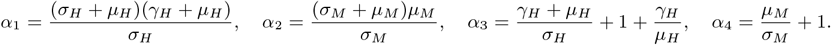

Then, lengthy algebraic computations show (details given in section S5 of the SM) that the between-host model has a unique endemic equilibrium

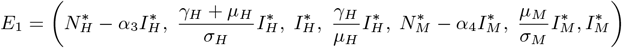

where the infected host class 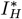 is

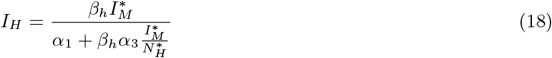

and the infected mosquito class 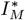 satisfies

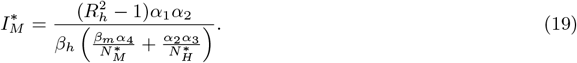

Note that 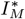 is positive if and only if the between-host reproduction number satisfies *R*_*h*_ *>* 1. Given the high dimensionality of the system, using standard linearization methods for *E*_1_ becomes analytically unfeasible. Hence, we use center manifold theory [14] to establish the local asymptotic stability of *E*_1_. To ease notation, we first relabel our variables as follows *x*_1_ = *E*_*H*_, *x*_2_ = *I*_*H*_, *x*_3_ = *R*_*H*_, *x*_4_ = *E*_*M*_, and *x*_5_ = *I*_*M*_. Let **f** = (*f*_1_, …, *f*_5_)^*T*^ be the vector field for the between-host model with 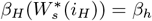 and *β*_*M*_ (*V* ^*^(*i*_*M*_)) = *β*_*m*_, such that 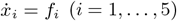, then

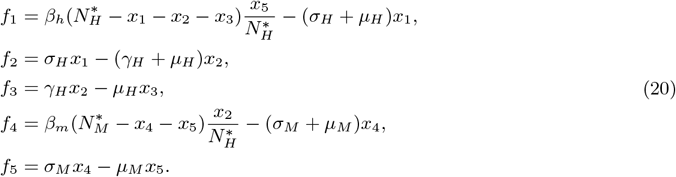

We are interested in the case where the between-host reproduction number satisfies *R*_*h*_ = 1. Furthermore, we will use *β*_*h*_ as a bifurcation parameter. From (17), the value of *β*_*h*_ such that *R*_*h*_ = 1 is

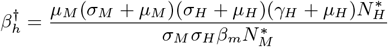

The linearization matrix of the transformed system (20) evaluated at the DFE when 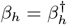 is

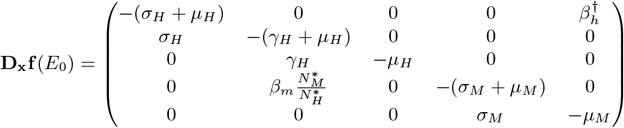

For the value 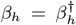, the matrix **D**_**x**_**f** (*E*_0_) has a simple zero eigenvalue, while all other eigenvalues are negative. Let **w** = (*w*_1_, …, *w*_5_)^*T*^ be the right eigenvector associated with the zero eigenvalue, then

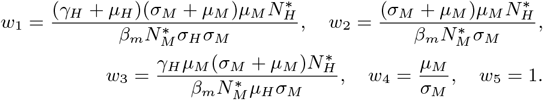

The left eigenvector of the zero eigenvalue, denoted as **v** = (*v*_1_, …, *v*_5_)^*T*^, is given by

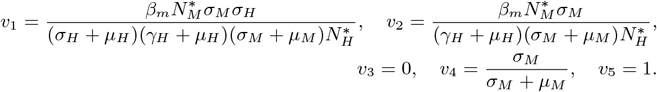

To determine the stability and direction of the bifurcation when *R*_*h*_ = 1, we calculate the coefficients *a* and *b*described in Theorem 4.1 in [14] and defined as

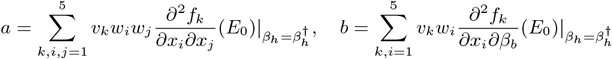

Algebraic calculations show that the only nonzero second partial derivatives with respect to the state variables of *f*_*k*_ (*k* = 1, …, 5) evaluated at the DFE are

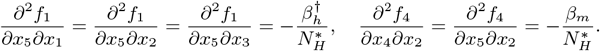

Since the above partial derivatives are negative and the left and right eigenvectors have nonnegative coordinates, it follows that *a <* 0. Likewise, a direct computation shows that

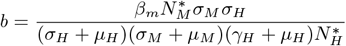

so *b >* 0. Hence, by Theorem 4.1 in [14], system (20) or, equivalently the uncoupled between-host model shows a forward bifurcation when *R*_*h*_ = 1 and the endemic equilibrium *E*_1_ is LAS when *R*_*h*_ *>* 1. These results are summarized in the next theorem.

#### Theorem 3.

*In the absence of feedback from the within-host and within-vector subsystems, if R*_*h*_ *<* 1, *then the DFE E*_0_ *is the only equilibrium point of the uncoupled between-host model* (1)*-*(2) *in* Ω *and is LAS. When R*_*h*_ *>* 1, *the model undergoes a forward transcritical bifurcation where the DFE becomes unstable, and there appears a unique endemic equilibrium E*_1_, *which is LAS*.

The theorem above indicates that the uncoupled between-host epidemic model exhibits classical threshold behavior, where *R*_*h*_ = 1 serves as the critical boundary between disease persistence and extinction. To identify the parameters that most influence these infected classes at the endemic equilibrium *E*_1_, we performed a GSA using Sobol’s method. Since host and vector populations are assumed to be at demographic equilibrium, only their ratio influences *R*_*h*_ and the overall system dynamics (see Section S3 in the SM). Accordingly, we normalize the total host population and vary the vector-to-host ratio by treating the mosquito mortality rate, *µ*_*M*_, as a free parameter, while keeping Λ_*H*_, *µ*_*H*_, and Λ_*M*_ fixed. The GSA results (Figure 6) indicate that *µ*_*M*_ is the dominant driver of infection prevalence in both mosquito and host populations across all three diseases. For infected hosts, the second most influential parameter is the intrinsic incubation period, 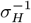. In the case of WNV, the transmission rate *β*_*H*_ also plays a notable role, with first- and total-order indices exceeding 0.2. For infected mosquitoes, the next most influential parameters are 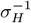 and *β*_*M*_, which have comparable effects but remain far less influential than *µ*_*M*_. These findings provide the foundation for the numerical exploration of the steady states of the coupled model (see Section 4.3).

**Figure 6:**
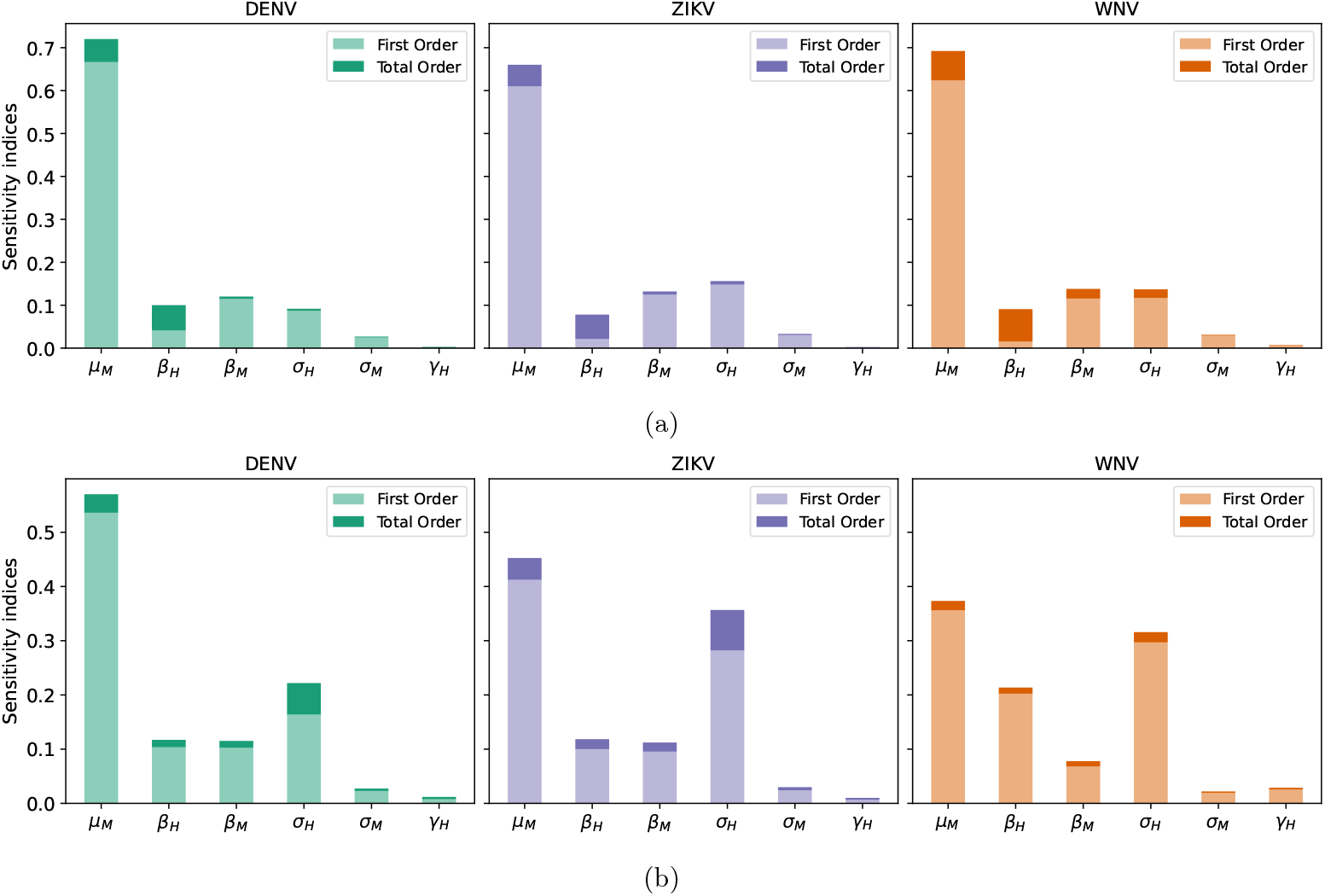
First and total order Sobol sensitivity indices for the infected mosquito (a) and host (b) classes at the equilibrium. The ranges used for the parameter values in the sensitivity analysis are based on 50% deviation from the mean values.

### 4.3 The coupled model

When the population-level dynamics are linked to the dynamics within the host and vector, with the fast system close to its unique steady state, the equilibrium values of the system are determined by the solution of the following system of equations (see section S5 of the SM):

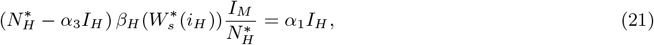

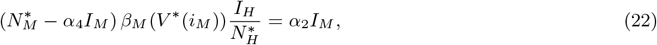

where 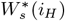 and *V* ^*^(*i*_*M*_) are given by (16) and (12), respectively. Observe that using equation (21) we can write *I*_*M*_ as a function of *I*_*H*_ as follows

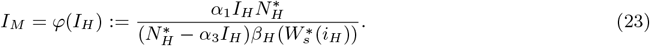

Likewise, using equation (22) we can express *I*_*H*_ as a function of *I*_*M*_ as follows

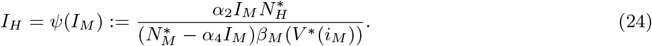

The equilibrium points of the coupled model are thus given by the intersection of the functions *φ*(*I*_*H*_) and *ψ*(*I*_*M*_). From the definition of *φ*(*I*_*H*_) and *ψ*(*I*_*M*_) it follows that the positive equilibrium points for the infected classes are bounded and satisfy 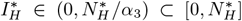 and 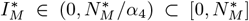.We start considering linear coupling functions 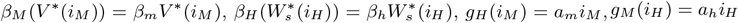. Under these assumptions, the within-vector equilibrium functions are

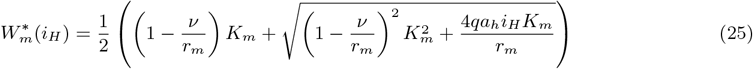

and

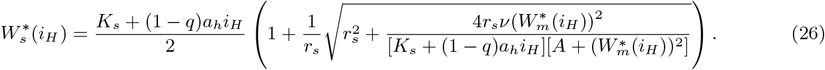

Therefore, *φ*(*I*_*H*_) becomes

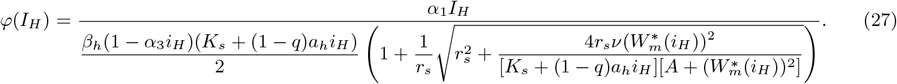

where 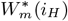 is defined in (25). Likewise, for *g*_*H*_ (*i*_*M*_) = *a*_*m*_*i*_*M*_, the function *ψ*(*I*_*M*_) becomes

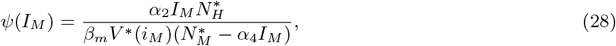

where the within-host equilibrium viral load is

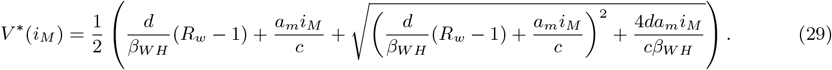

Since 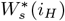 given by (26) is a monotonic increasing function of *i*_*H*_, the function *φ*(*I*_*H*_) given by (27) is a convex monotonic increasing function of *I*_*H*_ in the range 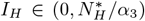.Likewise, since *V* ^*^(*i*_*M*_) given by (29) is a monotonic increasing function of *i*_*M*_, then *ψ*(*I*_*M*_) given by (28) is a convex monotonic increasing function of *I*_*M*_ in the range 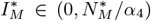. Therefore, for linear coupling functions, *φ*(*I*_*H*_) and *ψ*(*I*_*M*_) intersect at the origin and at most at one point with positive coordinates which leads to the following result.

#### Theorem 4.

*For linear coupling functions* 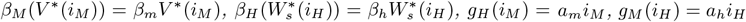, *the coupled multiscale model* (5) *presents always a disease-free equilibrium and at most one endemic equilibrium point*.

Theorem 4 implies that when using linear coupling transmission rates, the multiscale epidemic model (5) exhibits the same bifurcation structure as the uncoupled model. As shown in Section 3, the best fit to the viremia-infectiousness data from ZIKV-infected cynomolgus macaques and WNV-infected grackles follow a linear pattern (see also Suplemental Tables S4-S5). Figure 7 illustrates the functions *φ*(*I*_*H*_) (dashed lines) and *ψ*(*I*_*M*_) (solid lines), plotted using the mean parameter estimates for ZIKV under the linear coupling assumption. The corresponding functions for WNV display highly similar behavior and are therefore not shown. In all scenarios, the feedback terms are defined as *g*_*H*_ (*i*_*M*_) = *a*_*m*_*i*_*M*_ and *g*_*M*_ (*i*_*H*_) = *a*_*h*_*i*_*H*_, with the parameters set to *a*_*m*_ = 10^7^ and *a*_*h*_ = 10^7^. It is important to note that reliable empirical estimates for these feedback functions are currently unavailable, and the values used here are theoretical. Nonetheless, under these assumptions, both the within-host equilibrium viral load *V* ^*^(*i*_*M*_) and the virus concentration in the vector salivary glands 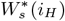 fall within biologically plausible ranges (refer to Figures 3b and 4a).

**Figure 7:**
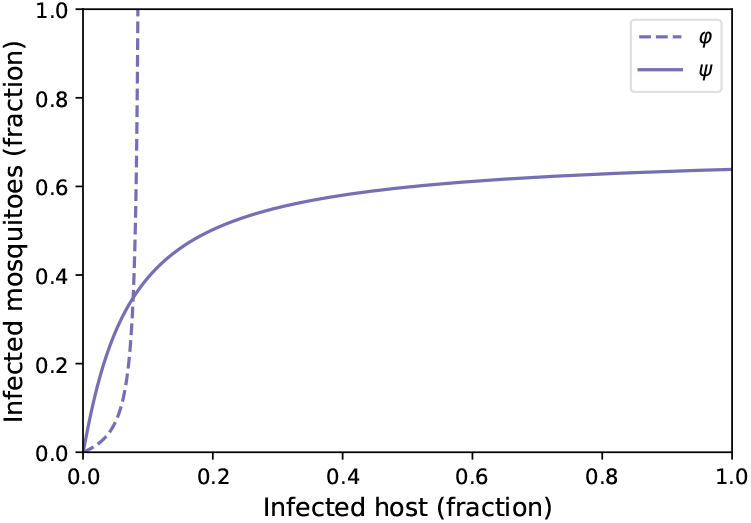
Functions *φ*(*I*_*H*_) (dashed lines) and *ψ*(*I*_*M*_) (solid lines) with the mean parameter values corresponding to ZIKV, and the linear coupling transmission rate derived from dose-response estimations using data from ZIKV-infected cynomolgus macaques. For all cases, *g*_*H*_ (*i*_*M*_) = *a*_*m*_*i*_*M*_, *g*_*M*_ (*i*_*H*_) = *a*_*h*_*i*_*H*_ with *a*_*m*_ = 10^7^, *a*_*h*_ = 10^7^.

Next, we investigate the dynamics of the coupled model for nonlinear coupling transmission functions. We focus on the Ferguson (7) and Hill (8) functions, both fitted to the viremia-infectiousness data in Figure 1. For DENV, both functions fit the data equally well based on AICc, indicating a sigmoidal relationship. Although the best fit model for the ZIKV and WNV data was the linear pattern, we also consider the estimates for the Ferguson and Hill functions to show how the model behaves under nonlinear assumptions. The sigmoidal dose-response models also performed reasonably well for these viruses; however, for WNV, the data covered only a narrow range of viremia values, limiting our ability to infer the relationship at low viremia levels.

Analytical determination of the equilibria in the coupled model becomes intractable when nonlinear coupling functions are employed. In particular, with nonlinear transmission rates, the convexity of the functions *φ*(*I*_*H*_) and *ψ*(*I*_*M*_) is no longer guaranteed. Since these functions depend on the equilibrium levels 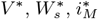, and 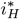, to investigate how the number of equilibrium points in the coupled system is influenced by changes in model parameters and the specific form of the transmission coupling functions, we perturbed the most influential parameters identified in the previous GSA: Λ_*T*_, *δ, p, c, K*_*s*_, *µ*_*M*_, and *σ*_*H*_. Each parameter was varied within *±*50% of its mean using a uniform distribution, across an ensemble of 2 samples. For each parameter set, we computed the number of intersections between *φ*(*I*_*H*_) and *ψ*(*I*_*M*_). The distribution of these intersection counts is illustrated in the histograms in Figure 8. Since *φ*(*I*_*H*_) and *ψ*(*I*_*M*_) always intersect at the disease-free equilibrium, the maximum number of positive (endemic) equilibrium points corresponds to the total number of intersections minus one.

**Figure 8:**
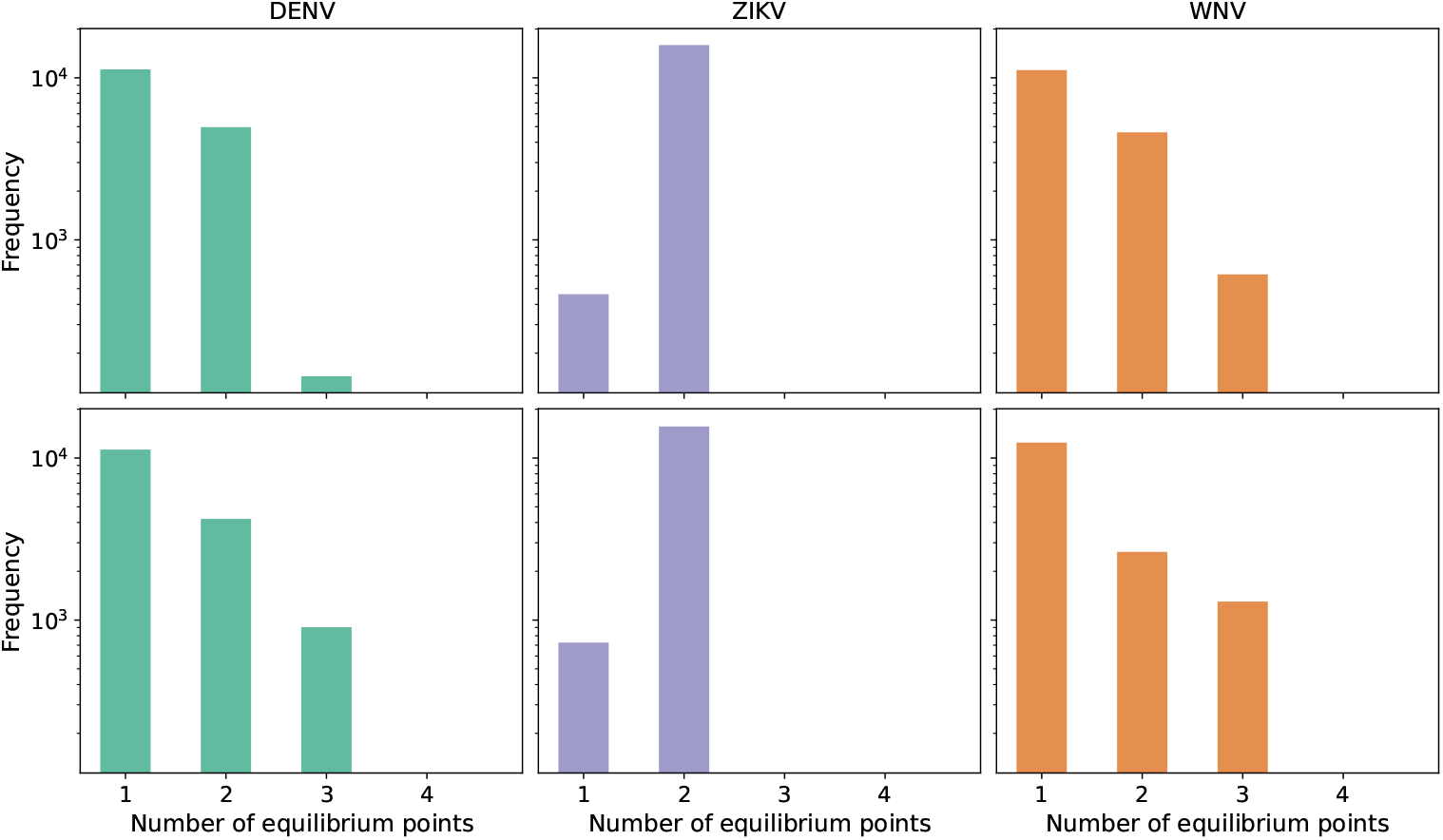
Histograms quantifying the number of times that the functions *φ*(*I*_*H*_) and *ψ*(*I*_*M*_) intersect (including the intersection at the DFE) for Ferguson (7) (top row), and Hill (8) (bottom row) coupling transmission functions and variations in the most influential parameters of the multiscale model. The x-axis gives the number of interceptions whereas the y-axis shows the frequency on a logarithmic scale. The first, second, and third columns show values corresponding to DENV, ZIKV, and WNV, respectively. For all cases, *g*_*H*_ (*i*_*M*_) = *a*_*m*_*i*_*M*_, *g*_*M*_ (*i*_*H*_) = *a*_*h*_*i*_*H*_ with parameters *a*_*m*_ = 10^7^, *a*_*h*_ = 10^7^.

Our results (see Figure 8) indicate that nonlinear dose-response relationships can give rise to multiple positive equilibrium points—unlike in linear scenarios, where such multiplicity does not occur. Specifically, in the cases of DENV and WNV, the model exhibits two positive equilibria under both the Ferguson and Hill functional forms. It is also worth noting that Figure 8 displays frequencies on a logarithmic scale to enhance visibility. While this representation may suggest that multiple positive equilibria are quite common, a linear-scale version (see Supplemental Figure S1) reveals that the actual frequency of a second positive equilibrium remains relatively low. Figure 9 shows an illustrative example of multiple crosses of the functions *φ*(*I*_*H*_) and *ψ*(*I*_*M*_) for the Hill-type transmission function and parameter values corresponding to DENV.

**Figure 9:**
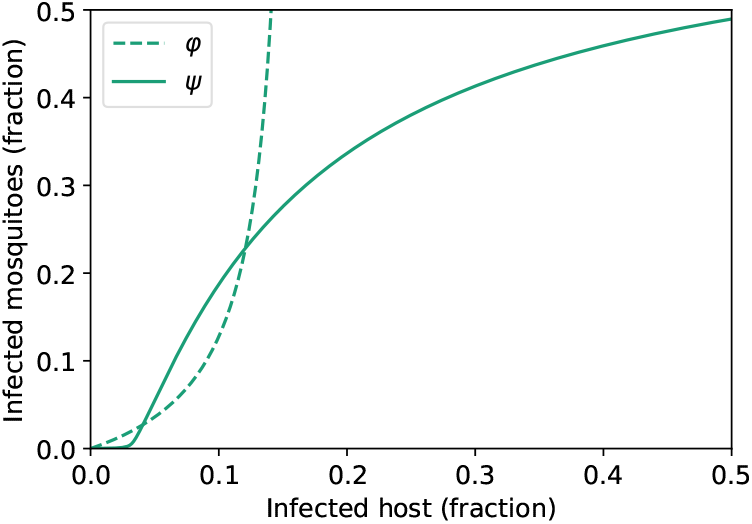
Functions *φ*(*I*_*H*_) (dashed line) and *ψ*(*I*_*M*_) (solid line) for the Hill-type transmission function and parameter values corresponding to DENV. The only parameter variation compared to the mean values is that the mean intrinsic incubation period 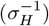 changes from 6 days to 4 days. The feedback functions are *g*_*H*_ (*i*_*M*_) = *a*_*m*_*i*_*M*_, *g*_*M*_ (*i*_*H*_) = *a*_*h*_*i*_*H*_ with *a*_*m*_ = 10^7^, *a*_*h*_ = 10^7^.

## 5 Discussion

Modeling has become an invaluable asset in epidemiology, enhancing our understanding of transmission dynamics and supporting the design of effective control strategies [24]. However, epidemiological models operating at the population level often neglect or simplify key biological processes occurring at finer scales, such as those within the host or the vector [36]. Multiscale modeling offers a pathway to investigate whether such simplification might compromise model accuracy or limit the relevance of outcomes, but must be pursued keeping in mind the importance of limiting complex implementations and untractable analysis.

In this study, we contribute to the multiscale epidemic modeling literature by proposing a new mechanistic modeling framework for mosquito-borne viruses explicitly connecting population-level host-vector transmission dynamics with within-host and within-vector viral processes. A distinctive feature of our framework is the incorporation of viral progression within the vector, particularly the transition of the virus from the midgut to the salivary glands, a process rarely modeled in previous multiscale models [38, 51]. Another significant feature is the bidirectional coupling between scales, allowing within-host processes to influence transmission rates and be reciprocally influenced by epidemiological dynamics. Prior progress in bidirectional scale-linking has largely focused on environmentally mediated diseases—typically using an additional equation to describe pathogen dynamics in a passive environmental reservoir [25, 26, 27, 72]. Our work extends this conceptual framework to vector-borne diseases where rather than treating the environment as a static intermediary, we consider within-vector viral kinetics in mosquitoes, which play an active role in pathogen transmission through host-seeking behavior.

The second contribution of this study is to assess the dynamic consequences of different functional forms of the transmission rate linking within-host and between-host processes. Our analytical results show that when the viremia-infectiousness relationship is linear, the multiscale model behaves qualitatively like the uncoupled model, retaining key features such as bifurcation structure. In such cases, incorporating an explicit multiscale framework adds unnecessary complexity, as the long-term behavior of the system can be adequately captured using a single-scale model. This suggests that the hierarchical multiscale structure is not crucial, meaning that the within-host scale can be studied independently, and key insights can be incorporated into the population-level model in a phenomenological way. In contrast, numerical simulations, using parameter values representative of Dengue virus (DENV), and West Nile virus (WNV), reveal that for nonlinear coupling transmission rates, the multiscale model may exhibit multiple positive equilibrium points and bifurcation structures that the uncoupled model does not capture. The existence of multiple steady states implies that epidemic outcomes can depend sensitively on initial conditions. In other words, small changes in initial number of infected individuals can result in vastly different prevalence levels at the endemic state. Moreover, the emergence of multiple endemic states also raises the possibility of a backward transcritical bifurcation, where the classic epidemiological threshold condition *R*_0_ *<* 1 no longer guarantees disease elimination. These multistability and bifurcation outcomes highlight the importance of implementing interventions early in the outbreak and also the need for more aggressive or sustained interventions (e.g., vaccination, quarantine) to effectively achieve disease eradication.

More broadly, this work contributes to the ongoing debate about when multiscale models are truly needed and when simpler, single-scale models are sufficient (see challenge 6 in Gog et al. [36]). Multiscale epidemic modeling introduces substantial complexity due to nonlinear feedbacks, a large number of parameters, and scarce empirical data. This complexity becomes even more pronounced in the context of vector-borne diseases, where it is necessary to explicitly model the vector’s role in pathogen transmission, adding another layer of interactions and parameters to the system. Hence, clarifying when multiscale approaches are necessary can guide more efficient and effective modeling choices [36, 56]. Our findings show that the careful empirical validation of the dose-response relationship linking within-host viral load to transmission rates is one of the ways to distinguish between a case where the multiscale framework is needed (nonlinear functions) or not (linear functions). However, in the absence of empirical data, modelers often default to assuming a linear function for parsimony. This assumption may not be biologically justified and could undermine the potential advantages of a multiscale approach.

A key strength of our modeling approach lies in the integration of empirical dose-response data into the transmission function, which contributes to the biological validation of the model. Specifically, we gathered existing datasets that associate infectious viral titers in vertebrate hosts with the likelihood of mosquito infection for DENV, ZIKV, and WNV. These data were exclusively sourced from studies in which mosquitoes fed directly on live hosts, offering a more realistic representation of natural transmission scenarios. We note that scientists performing such experiments might not be aware that their datasets could greatly contribute to improving epidemiological models’ accuracy. Efforts should be done to enhance the dialogue between these disciplines and improve the availability and reusability of similar datasets.

Another advantage of our model is that it can be easily extendable to include more compartments, and hence, it can be adapted to represent different vector-host-pathogen systems. At the within-host level, the model can be expanded to incorporate immune responses explicitly. The within-vector model can be further refined to capture the infection, dissemination, and transmission barriers more accurately [51]. Still, we remind that this study does not aim to produce quantitative predictions of disease spread. Instead, we aim to introduce a theoretical framework to explore how the functional form of the transmission rate linking within-host and between-host processes, shapes long-term, population-level dynamics.

Keeping the multiscale epidemic model both simple and manageable unavoidably results in certain limitations. For instance, our analysis relies on time-scale separation methods, which overlook the effects of transient within-host and within-vector dynamics at the population level [5, 33, 43]. The model also assumes uniform immunological dynamics among vertebrate hosts and vectors, neglecting individual variability at the within-host scale. In other words, the within-host model represents the average dynamics of a typical host. This limitation could be addressed by employing a time-since-infection modeling framework using partial differential equations e.g. [13, 38, 44, 55], or by indexing within-host variables by individual to quantify the aggregate viral load across the population [25]. However, such approaches would significantly increase mathematical complexity. An alternative path would be to develop an individual-based multiscale model e.g. [49, 70], linking within-host processes to network-based epidemiological models. While computationally tractable, such approaches often provide limited theoretical insight and require extensive parameterization and empirical data for validation. Another limitation of the model is that the influence of the within-host scale is restricted to transmission rates, while other parameters, such as recovery or mortality, may also depend on intra-host dynamics [18, 56, 58]. We focused on transmission in this study due to the availability of empirical viremia-infectiousness data, but future work should explore whether our key findings hold when virulence and recovery rates are also modeled as functions of within-host variables. Despite these limitations, the proposed model captures the key features of mosquito-borne diseases and serves as a foundation for future development of more detailed multiscale models.

## Data availability

The datasets analyzed during the current study are publicly available within the references cited in the article. No new datasets were generated or analyzed during the current study. Relevant code will be uploaded to a GitLab repository once the paper has been accepted for publication.

## Competing interests

The authors declare they have no known competing financial interests that could have appeared to influence the results reported in this work.

## Funding

Funded by the European Union (WiLiMan-ID, grant agreement 101083833). Views and opinions expressed are however those of the authors only and do not necessarily reflect those of the European Union or REA. Neither the European Union nor the granting authority can be held responsible for them. This work also received the support of the French region Pays de la Loire.

## Acknowledgments

We are grateful to the WiLiMan-ID consortium for the valuable discussions held during the second annual meeting in Vienna. We extend special thanks to Dr. Quirine ten Bosch for her insightful contributions.

## CRediT authorship contribution statement

**Fernando Saldaña:** Conceptualization, Methodology, Formal analysis, Software, Visualization, Writing – original draft, Writing – review and editing. **Jorge X. Velasco-Hernández:** Conceptualization, Formal analysis, Writing – review and editing. **Pauline Ezanno:** Conceptualization, Supervision, Funding Acquisition, Project Administration, Writing – review and editing. **Hélène Cecilia:** Conceptualization, Software, Visualization, Supervision, Writing - original draft, Writing – review and editing.

## Notes

### Competing Interest Statement

The authors have declared no competing interest.

