## Supplementary material for "Multiscale Modeling of Vector-Borne Diseases: The Role of Dose-Dependent Transmission": Suplementary Material

#### S1 Model parameters

The mean parameter values for the between-host model, along with their sources, are summarized in Table S1. We consider Dengue virus (DENV), Zika virus (ZIKV), and West Nile virus (WNV) as representative examples of major mosquito-borne viruses. For DENV and ZIKV, we retrieved values from studies focusing on humans as vertebrate hosts, while the vectors are *Aedes aegypti* and *Aedes albopictus* mosquitoes. For DENV the parameter values were estimated using single serotype models so they represent an average of the four DENV serotypes. For WNV, the data correspond to *Culex pipiens* mosquitoes as the vector, with House sparrows (*Passer domesticus*) and American robins (*Turdus migratorius*) as the vertebrate hosts, based on studies from [11] and [13]. To simulate the early stage of an outbreak, the initial conditions assume that 1% of both host and mosquito populations are infected, with the remaining individuals classified as susceptible.

Table S1: Parameters of the population-level epidemic model. The time unit is days. The vector-to-host ratio is fixed based on $R_h = 2$. Par. = parameter

| **Par.** | **Description** | **Virus** | **Value** | **Units** | **Ref** |
|---|---|---|---|---|---|
| $\mu_M$ | Mosquito death rate | DENV | 0.07 | day$^{-1}$ | [2] |
| | | ZIKV | 0.066 | day$^{-1}$ | [12] |
| | | WNV | 0.12 | day$^{-1}$ | [11] |
| $\beta_h$ | Mosquito-to-host transmission rate | DENV | 0.67 | day$^{-1}$ | [14] |
| | | ZIKV | 0.96 | day$^{-1}$ | [12] |
| | | WNV | 0.44 | day$^{-1}$ | [13] |
| $\beta_m$ | Host-to-mosquito transmission rate | DENV | 0.39 | day$^{-1}$ | [14] |
| | | ZIKV | 0.26 | day$^{-1}$ | [12] |
| | | WNV | 0.974 | day$^{-1}$ | [13] |
| $\sigma_H^{-1}$ | Intrinsic incubation period | DENV | 6 | day | [2] |
| | | ZIKV | 8.5 | day | [12] |
| | | WNV | 0.5 | day | [11] |
| $\sigma_M^{-1}$ | Extrinsic incubation period | DENV | 8 | day | [2] |
| | | ZIKV | 7 | day | [2, 12] |
| | | WNV | 3.5 | day | [11] |
| $\gamma_H^{-1}$ | Hosts infectious period | DENV | 4.5 | day | [14] |
| | | ZIKV | 5.5 | day | [14] |
| | | WNV | 2.3 | day | [11] |
| $N_M^*/N_H^*$ | Vector-to-host ratio | DENV | 14.42 | | Assumed |
| | | ZIKV | 17.42 | | Assumed |
| | | WNV | 3.5 | | Assumed |

Since both host and vector populations are in demographic equilibrium, it is their ratio —not their absolute sizes— that affects $R_h$ and the overall model dynamics. This allows us to normalize the populations. Specifically, we assume $\Lambda_H = \mu_H = 1$, resulting in a host population of $N_H^* = 1$. Likewise, the vector recruitment rate $\Lambda_M$ can be fixed allowing the mosquito death rate $\mu_M$ to serve as a free parameter for controlling the vector-to-host ratio. We choose $\Lambda_M/\mu_M$ based on reasonable ranges for the between-host reproduction number $R_h$ (see Figure 5 in the main text). In particular, we fixed $R_h = 2$ as a mean value.

Table S2: Parameters values of the within-host model. The viral load is measured in plaque-forming units per milliliter (PFU/ml). Note that for the death rate $d$, we found values only for DENV and, for simplicity, applied the same value to ZIKV and WNV. The production rate of susceptible target cells $\Lambda_T$ (and $\beta_{WH}$ for DENV) were selected fixing $R_w = 3$. Par. = parameter

| Par. | Description | Virus | Value | Units | Ref |
|---|---|---|---|---|---|
| $d$ | Death rate of susceptible target cells | DENV | 0.03 | day$^{-1}$ | [1] |
| | | ZIKV | 0.03 | day$^{-1}$ | Assumed |
| | | WNV | 0.03 | day$^{-1}$ | Assumed |
| $\Lambda_T$ | Production rate susceptible target cells | DENV | 590 | cells/day | Assumed |
| | | ZIKV | 159 | cells/day | Assumed |
| | | WNV | 4020 | cells/day | Assumed |
| $\beta_{WH}$ | Within-host infection rate | DENV | $1 \times 10^{-4}$ | ml/day | Assumed |
| | | ZIKV | $2 \times 10^{-5}$ | ml/day | [5] |
| | | WNV | $4 \times 10^{-4}$ | ml/day | [3] |
| $k^{-1}$ | Duration of the eclipse phase | DENV | 0.25 | day | [4, 7] |
| | | ZIKV | 0.25 | day | [5] |
| | | WNV | 0.33 | day | [3] |
| $\delta$ | Death rate of infected target cells | DENV | 2.5 | day$^{-1}$ | [1] |
| | | ZIKV | 3.5 | day$^{-1}$ | [5] |
| | | WNV | 23.03 | day$^{-1}$ | [3] |
| $p$ | In-host viral production rate per cell | DENV | 50 | PFU/(cell day) | [1] |
| | | ZIKV | 1000 | PFU/(cell day) | [5] |
| | | WNV | 57.88 | PFU/(cell day) | [3] |
| $c$ | In-host clearance rate of free virions | DENV | 13 | day$^{-1}$ | [1] |
| | | ZIKV | 10 | day$^{-1}$ | [5] |
| | | WNV | 44.43 | day$^{-1}$ | [3] |

The parameter values used in the within-host model are summarized in Table S2. For the death rate $d$, we found values only for DENV and, for consistency, applied the same value to ZIKV and WNV. Similarly, we could not find data for the production rate of susceptible target cells $\Lambda_T$. To allow for comparison between diseases, we chose $\Lambda_T$ such that the within-host reproduction number $R_w$ for each virus is equal to 3, while also ensuring the model captures typical features of arbovirus infections. For the case of DENV, we also selected $\beta_{WH}$ following the same justification. Certainly, since these values are not derived from empirical data, this represents a limitation of the study. The initial susceptible target cell density $T(0)$ is $8 \times 10^4$ for DENV [4], $1 \times 10^4$ for ZIKV [5], and $2.3 \times 10^5$ for WNV [3]. For simplicity, the initial density of virus $V(0)$ is assumed equal to 1 PFU/ml across all three viruses. Likewise, the initial density of exposed and infected cells is assumed equal to zero.

Table S3: Parameters values of the within-vector model. The time unit is days, and the viral load is measured in PFU/ml. Note that in the original data source viral load is measured in $\text{TCDI}_{50}$/ml but in the simulations we use a conversion factor from [6] to obtain PFU to be consistent with the rest of parameter values. Par. = parameter

| Par. | Description | Value | Units | Ref |
|---|---|---|---|---|
| $r_m$ | Viral growth rate in the midgut | 1.26 | day$^{-1}$ | [15] |
| $K_m$ | Viral carrying capacity in the midgut | $4.33 \times 10^5$ | $\text{TCDI}_{50}$/ml | [15] |
| $\nu$ | Viral dissemination rate | 0.11 | day$^{-1}$ | [15] |
| $A$ | Half saturation constant | $1.95 \times 10^4$ | $\text{TCDI}_{50}{}^2$/ml$^2$ | [15] |
| $r_s$ | Viral growth rate in the salivary glands | 1.77 | day$^{-1}$ | [15] |
| $K_s$ | Viral carrying capacity in the salivary glands | $6.84 \times 10^6$ | $\text{TCDI}_{50}$/ml | [15] |

For the within-vector model, parameter values were available only for *Aedes aegypti* mosquitoes infected with ZIKV, as summarized in Table S3. Following the approach in [15], the initial viral load in the mosquito's midgut $W_m(0)$ is set to 3 PFU/ml, while the salivary glands start with a viral load $W_s(0)$ of 0, reflecting that the virus should first cross the dissemination barrier before reaching the salivary glands.

### S2 Dose-response relationships

Three functional forms (Eqs. 6–8 in the main text) were evaluated to characterize the dose–response relationship between infectious titers in vertebrate hosts and the probability of mosquito infection, using a maximum likelihood estimation framework. Mosquito infection was assessed by viral detection either in the body (abdomen excluding head, legs, and wings) or in the legs, depending on the dataset. The vertebrate hosts employed in the dose–response experiments were humans for DENV, cynomolgus macaques for ZIKV, and common grackles for WNV.

Each functional form was fitted using either a binomial or betabinomial likelihood, with the latter applied to account for overdispersion. Model performance was compared using the corrected Akaike Information Criterion (AICc; Table S4). Across all datasets, the betabinomial likelihood consistently outperformed the binomial, indicating overdispersion. For ZIKV and WNV, the linear model provided the best fit. For DENV, both the Ferguson and Hill models performed equally well according to AICc. To quantify uncertainty in the fitted curves, $1,000$ parameter sets were sampled from a multivariate normal distribution defined by the estimated covariance matrix obtained during the fitting process. Point estimates and corresponding 95% confidence intervals for each parameter are reported in Table S5.

Table S4: Model selection for dose-response relationships predicting mosquito infection probabilities based on vertebrate host infectious titers. The table reports corrected Akaike Information Criterion (AICc); models with the lowest AICc (in bold) are preferred. Equation numbers refer to the dose-response functional forms described in the main text. df = degrees of freedom.

| **Dose-response functional form** | **Likelihood** | **df** | **AICc** | | |
|---|---|---|---|---|---|
| | | | **DENV** | **ZIKV** | **WNV** |
| linear (Eq. 6) | binomial | 1 | 264.0 | 50.4 | 102.7 |
| | beta-binomial | 2 | 104.1 | **45.1** | **82.3** |
| Ferguson (Eq. 7) | binomial | 2 | 164.8 | 48.2 | 103.6 |
| | beta-binomial | 3 | **88.0** | 46.9 | 85.2 |
| Hill (Eq. 8) | binomial | 2 | 166.7 | 54.4 | 103.6 |
| | beta-binomial | 3 | **88.0** | 49.9 | 85.2 |

Table S5: Parameter estimates for the dose-response functional forms (Eqs. 6–8 in the main text). Par. = parameter, od = overdispersion.

| Virus | Data source | Functional form | Mosquito infection | Par. | Point estimate | 95% CI |
|---|---|---|---|---|---|---|
| DENV-4 | [9] | linear | Legs | $a$ | 0.074 | [0.051 ; 0.11] |
| | | | | od | 0.60 | [0.30 ; 1.20] |
| DENV-4 | [9] | Ferguson | Legs | $\theta_0$ | 6.12 | [5.73 ; 6.52] |
| | | | | $\theta_1$ | 6.77 | [4.20 ; 10.92] |
| | | | | od | 1.24 | [0.56 ; 2.74] |
| DENV-4 | [9] | Hill | Legs | $\gamma_0$ | $e^{16}$ | $[e^{6.99} ; e^{25.03}]$ |
| | | | | $\gamma_1$ | 9.20 | [5.24 ; 16.16] |
| | | | | od | 1.21 | [0.55 ; 2.66] |
| ZIKV | [10] | linear | Legs | $a$ | 0.16 | [0.13 ; 0.20] |
| | | | | od | 4.45 | [1.16 ; 17.13] |
| ZIKV | [10] | Ferguson | Legs | $\theta_0$ | 4.19 | [3.81 ; 4.60] |
| | | | | $\theta_1$ | 4.37 | [2.25 ; 8.48] |
| | | | | od | 4.45 | [1.30 ; 24.74] |
| ZIKV | [10] | Hill | Legs | $\gamma_0$ | $e^{6.69}$ | $[e^{-0.38} ; e^{13.77}]$ |
| | | | | $\gamma_1$ | 5.13 | [1.98 ; 13.29] |
| | | | | od | 5.95 | [1.19 ; 29.81] |
| WNV | [17] | linear | Bodies | $a$ | 0.066 | [0.053 ; 0.084] |
| | | | | od | 8.88 | [2.96 ; 26.58] |
| WNV | [17] | Ferguson | Bodies | $\theta_0$ | 7.91 | [6.20 ; 10.09] |
| | | | | $\theta_1$ | 2.93 | [0.91 ; 9.42] |
| | | | | od | 9.60 | [3.19 ; 28.86] |
| WNV | [17] | Hill | Bodies | $\gamma_0$ | $e^{7.96}$ | $[e^{-1.55} ; e^{17.48}]$ |
| | | | | $\gamma_1$ | 4.11 | [1.19 ; 14.19] |
| | | | | od | 9.63 | [3.20 ; 28.92] |

### S3 Time-scale separation

The following time-scale separation allows us to conduct the fast-slow analysis presented in the main text. For the convenience of the reader, we present the four subsystems comprising the full multiscale model, namely, the vector-to-host transmission model

$$\begin{aligned}\frac{dS_H}{dt} &= \Lambda_H - \beta_H(W_s)S_H\frac{I_M}{N_H} - \mu_H S_H,\\ \frac{dE_H}{dt} &= \beta_H(W_s)S_H\frac{I_M}{N_H} - (\sigma_H + \mu_H)E_H,\\ \frac{dI_H}{dt} &= \sigma_H E_H - (\gamma_H + \mu_H)I_H,\\ \frac{dR_H}{dt} &= \gamma_H I_H - \mu_H R_H\end{aligned} \tag{S1}$$

the host-to-vector transmission model

$$\begin{aligned}\frac{dS_M}{dt} &= \Lambda_M - \beta_M(V)S_M\frac{I_H}{N_H} - \mu_M S_M,\\ \frac{dE_M}{dt} &= \beta_M(V)S_M\frac{I_H}{N_H} - (\sigma_M + \mu_M)E_M,\\ \frac{dI_M}{dt} &= \sigma_M E_M - \mu_M I_M,\end{aligned} \tag{S2}$$

the within-host model

$$\begin{aligned}\frac{dT}{d\tau} =& \Lambda_T - \beta_{WH}TV - dT,\\ \frac{dL}{d\tau} =& \beta_{WH}TV - (k+d)L,\\ \frac{dY}{d\tau} =& kL - \delta Y,\\ \frac{dV}{d\tau} =& g_H(i_M) + pY - cV,\end{aligned} \tag{S3}$$

and the within-vector model

$$\begin{aligned}\frac{dW_m}{d\tau} =& qg_M(i_H) + r_m W_m\left(1 - \frac{W_m}{K_m}\right) - \nu W_m,\\ \frac{dW_s}{d\tau} =& \frac{\nu W_m^2}{A + W_m^2} + r_s W_s\left(1 - \frac{W_s}{K_s + (1-q)g_M(i_H)}\right).\end{aligned} \tag{S4}$$

If we assume that the slow and fast time scales are related as $t = \epsilon\tau$ where $0 < \epsilon \ll 1$, then the chain rule allows us to rewrite the slow subsystems in terms of the fast scale as:

$$\begin{aligned}\frac{dS_H}{d\tau} =& \epsilon\left(\Lambda_H - \beta_H(W_s)S_H\frac{I_M}{N_H} - \mu_H S_H\right),\\ \frac{dE_H}{d\tau} =& \epsilon\left(\beta_H(W_s)S_H\frac{I_M}{N_H} - (\sigma_H + \mu_H)E_H\right),\\ \frac{dI_H}{d\tau} =& \epsilon\left(\sigma_H E_H - (\gamma_H + \mu_H)I_H\right),\\ \frac{dR_H}{d\tau} =& \epsilon\left(\gamma_H I_H - \mu_H R_H\right),\\ \frac{dS_M}{d\tau} =& \epsilon\left(\Lambda_M - \beta_M(V)S_M\frac{I_H}{N_H} - \mu_M S_M\right),\\ \frac{dE_M}{d\tau} =& \epsilon\left(\beta_M(V)S_M\frac{I_H}{N_H} - (\sigma_M + \mu_M)E_M\right),\\ \frac{dI_M}{d\tau} =& \epsilon\left(\sigma_M E_M - \mu_M I_M\right).\end{aligned} \tag{S5}$$

Hence, the full multiscale model in terms of the scale $\tau$ is given by subsystems (S5), (S3), and (S4). The fast dynamics can be analyzed by letting $\epsilon = 0$. This implies that the derivative of the vector of slow state variables $x = (S_H, E_H, I_H, R_H, S_M, E_M, I_M)^T$ is equal to zero on the scale $\tau$. Hence, we can investigate the fast dynamics, assuming that the slow state variables are constant.

Likewise, we can rewrite the fast within-host subsystems in terms of the slow time scale $t$ as:

$$\begin{aligned}\epsilon\frac{dT}{dt} =& \Lambda_T - \beta_{WH}TV - dT,\\ \epsilon\frac{dL}{dt} =& \beta_{WH}TV - (k+d)L,\\ \epsilon\frac{dY}{dt} =& kL - \delta Y,\\ \epsilon\frac{dV}{dt} =& g_H(i_M) + pY - cV,\\ \epsilon\frac{dW_m}{dt} =& qg_M(i_H) + r_m W_m\left(1 - \frac{W_m}{K_m}\right) - \nu W_m,\\ \epsilon\frac{dW_s}{dt} =& \frac{\nu W_m^2}{A + W_m^2} + r_s W_s\left(1 - \frac{W_s}{K_s + (1-q)g_M(i_H)}\right).\end{aligned} \tag{S6}$$

Therefore, the multiscale model in terms of the scale $t$ is composed of subsystems (S1), (S2), and (S6), and we can analyze the slow dynamics assuming $\epsilon = 0$ in these subsystems. Setting $\epsilon = 0$ in (S6) implies that the fast system can be treated as being in a steady state on the scale $t$.

### S4 Computation of the basic reproduction number via the next-generation method

We obtain the basic reproduction numbers using the next-generation matrix [8] and the method of van den Driessche and Watmough [16]. In general terms, to compute the next-generation matrix

$\mathbf{K}$, it is necessary to determine the subsystem that describes the production of new infections and changes in state among the infected classes. The Jacobian matrix $\mathbf{J}$ corresponding to the linearization of this subsystem at the disease-free equilibrium is decomposed as $\mathbf{F}-\mathbf{V}$. The matrix $\mathbf{F}$ is the transmission part, describing the production of new infections, and $\mathbf{V}$ describes changes in status, such as recovery or death. Then, the next-generation matrix is given by $\mathbf{K} = \mathbf{F}\mathbf{V}^{-1}$, and the basic reproduction number is defined as the spectral radius of $\mathbf{K}$.

We first compute the reproduction number, denoted $R_w$, for the within-host model (S3) with $g_H(i_M) = 0$. Let $L$, $Y$, and $V$ represent the infected compartments. The matrices $\mathbf{F}$ and $\mathbf{V}$, which correspond to the new infection terms and the remaining transfer terms, are defined as follows:

$$\mathbf{F} = \begin{pmatrix} 0 & 0 & \beta_{WH}\hat{T} \\ 0 & 0 & 0 \\ 0 & 0 & 0 \end{pmatrix}, \quad \mathbf{V} = \begin{pmatrix} k+d & 0 & 0 \\ -k & \delta & 0 \\ 0 & -p & c \end{pmatrix}.$$

The inverse of $\mathbf{V}$ can be easily obtained as

$$\mathbf{V}^{-1} = \begin{pmatrix} \dfrac{1}{k+d} & 0 & 0 \\ \dfrac{k}{\delta(k+d)} & \dfrac{1}{\delta} & 0 \\ \dfrac{kp}{\delta c(k+d)} & \dfrac{p}{\delta c} & \dfrac{1}{c} \end{pmatrix}.$$

Hence, the next-generation matrix is given by

$$\mathbf{K} = \mathbf{F}\mathbf{V}^{-1} = \begin{pmatrix} \dfrac{pk\beta_{WH}\hat{T}}{c(k+d)\delta} & \dfrac{p\beta_{WH}\hat{T}}{c\delta} & \dfrac{\beta_{WH}\hat{T}}{c} \\ 0 & 0 & 0 \\ 0 & 0 & 0 \end{pmatrix}.$$

Observe that $\mathbf{K}$ has two eigenvalues equal to zero and one positive eigenvalue equal to $\frac{pk\beta_{WH}\hat{T}}{c(k+d)\delta}$. The nonzero eigenvalue of $\mathbf{K}$ thus gives the following analytical expression for $R_w$:

$$R_w = \frac{p}{\delta}\frac{1}{c}\beta_{WH}\hat{T}\frac{k}{k+d}.$$

For the between-host epidemic system with constant transmission rates $\beta_H(W_s) = \beta_h$ and $\beta_M(V) = \beta_m$, we obtain the following next-generation matrix

$$\hat{\mathbf{K}} = \begin{pmatrix} 0 & 0 & \dfrac{\sigma_M\beta_h}{(\sigma_M+\mu_M)\mu_M} & \dfrac{\beta_h}{\mu_M} \\ 0 & 0 & 0 & 0 \\ \dfrac{\sigma_H\beta_m N_M^*}{(\sigma_H+\mu_H)(\gamma_H+\mu_H)N_H^*} & \dfrac{\beta_m N_M^*}{(\gamma_H+\mu_H)N_H^*} & 0 & 0 \\ 0 & 0 & 0 & 0 \end{pmatrix}$$

Clearly, $rank(\hat{\mathbf{K}}) = 2$ and the two nonzero eigenvalues of $\hat{\mathbf{K}}$ are given by

$$\lambda_{+,-} = \pm\sqrt{\frac{\sigma_M\beta_h}{(\sigma_M+\mu_M)\mu_M} \times \frac{\sigma_H\beta_m N_M^*}{(\sigma_H+\mu_H)(\gamma_H+\mu_H)N_H^*}}.$$

Therefore $R_h = \lambda_+$.

### S5 Equilibrium solutions of the between-host model

The solution of the following system gives the equilibrium points of the uncoupled between-host model:

$$\begin{aligned} 0 &= \beta_h(N_H^* - E_H - I_H - R_H)\frac{I_M}{N_H^*} - (\sigma_H+\mu_H)E_H, \\ 0 &= \sigma_H E_H - (\gamma_H+\mu_H)I_H, \\ 0 &= \gamma_H I_H - \mu_H R_H, \\ 0 &= \beta_m(N_M^* - E_M - I_M)\frac{I_H}{N_H^*} - (\sigma_M+\mu_M)E_M, \\ 0 &= \sigma_M E_M - \mu_M I_M. \end{aligned} \tag{S7}$$

Notice that the equations for the susceptible classes were omitted because $S_H = N_H^* - E_H - I_H - R_H$ and $S_M = N_M^* - E_M - I_M$. From (S7), we obtain

$$E_H = \frac{(\gamma_H + \mu_H)I_H}{\sigma_H}, \quad R_H = \frac{\gamma_H I_H}{\mu_H}, \quad E_M = \frac{\mu_M I_M}{\sigma_M}. \tag{S8}$$

Substituting (S8) into the first and fourth equations of the system (S7), we have

$$\beta_h \left(N_H^* - \alpha_3 I_H\right) \frac{I_M}{N_H^*} = \alpha_1 I_H, \tag{S9}$$

$$\beta_m \left(N_M^* - \alpha_4 I_M\right) \frac{I_H}{N_H^*} = \alpha_2 I_M, \tag{S10}$$

where

$$\alpha_1 = \frac{(\sigma_H + \mu_H)(\gamma_H + \mu_H)}{\sigma_H}, \quad \alpha_2 = \frac{(\sigma_M + \mu_M)\mu_M}{\sigma_M}, \quad \alpha_3 = \frac{\gamma_H + \mu_H}{\sigma_H} + 1 + \frac{\gamma_H}{\mu_H} \quad \alpha_4 = \frac{\mu_M}{\sigma_M} + 1.$$

From (S9) we can express the class $I_H$ in terms of the class $I_M$ as follows:

$$I_H = \frac{\beta_h I_M}{\alpha_1 + \beta_h \alpha_3 \frac{I_M}{N_H^*}} \tag{S11}$$

Substituting (S11) into (S10) gives

$$\beta_m \beta_h \frac{N_M^*}{N_H^*} I_M - \frac{\beta_m \alpha_4 \beta_h}{N_M^*} I_M^2 = \alpha_1 \alpha_2 I_M + \frac{\beta_h \alpha_2 \alpha_3}{N_H^*} I_M^2$$

which, after simplification, yields

$$\beta_h \left( \frac{\beta_m \alpha_4}{N_M^*} + \frac{\alpha_2 \alpha_3}{N_H^*} \right) I_M^2 + (1 - R_h^2)\alpha_1 \alpha_2 I_M = 0. \tag{S12}$$

The roots of the above quadratic polynomial determine the equilibrium values. Clearly $I_M = 0$ is always a solution of (S12), therefore the between-host model always has a disease-free equilibrium $E_0 = (N_H^*, 0, 0, 0, N_M^*, 0, 0)$. The positive root of (S12) gives

$$I_M^* = \frac{(R_h^2 - 1)\alpha_1 \alpha_2}{\beta_h \left( \frac{\beta_m \alpha_4}{N_M^*} + \frac{\alpha_2 \alpha_3}{N_H^*} \right)}, \tag{S13}$$

which is positive if and only if the between-host reproduction number satisfies $R_h > 1$. Substituting (S13) into (S11), we obtain the infected host class at the endemic equilibrium $I_H^*$, and hence we have proven the existence of a unique endemic equilibrium point whose coordinates are

$$E_1 = \left( N_H^* - \alpha_3 I_H^*, \; \frac{\gamma_H + \mu_H}{\sigma_H} I_H^*, \; I_H^*, \; \frac{\gamma_H}{\mu_H} I_H^*, \; N_M^* - \alpha_4 I_M^*, \; \frac{\mu_M}{\sigma_M} I_M^*, I_M^* \right)$$

This analysis implies that when the between-host and within-host models are coupled, with the fast systems close to its unique steady state, the equilibrium values of the system are determined by the solution of the following system:

$$\beta_H(W_s^*(i_H)) \left(N_H^* - \alpha_3 I_H\right) \frac{I_M}{N_H^*} = \alpha_1 I_H,$$

$$\beta_M(V^*(i_M)) \left(N_M^* - \alpha_4 I_M\right) \frac{I_H}{N_H^*} = \alpha_2 I_M.$$

### S6 Supplementary figures

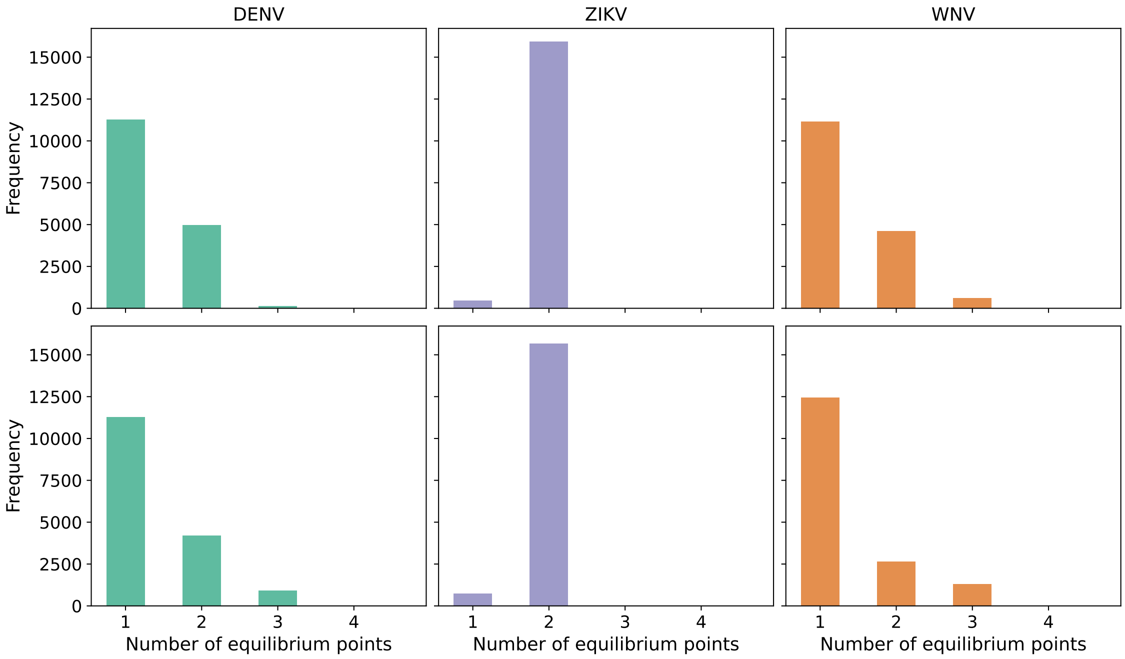


Figure S1: Histograms quantifying the number of times that the functions $\varphi(I_H)$ and $\psi(I_M)$ intersect (including the intersection at the DFE) for Ferguson (top row), and Hill (bottom row) coupling transmission functions and variations in the most influential parameters of the multiscale model. The x-axis gives the number of interceptions whereas the y-axis shows the frequency in linear scale. The first, second, and third columns show values corresponding to DENV, ZIKV, and WNV, respectively. For all cases, $g_H(i_M) = a_m i_M$, $g_M(i_H) = a_h i_H$ with parameters $a_m = 10^7$, $a_h = 10^7$.
